# Calreticulin is required for cuticle deposition and trabeculae formation inside butterfly wing scale cells

**DOI:** 10.1101/2024.09.12.612608

**Authors:** Ru Hong, Cédric Finet, Antónia Monteiro

## Abstract

Insect cuticle is normally deposited outside the plasma membrane of epidermal cells, making it unclear how cuticular pillars (trabeculae) are found inside butterfly wing scale cells. By co-labelling the cuticle and the plasma membrane, we found evidence that the plasma membrane invaginates towards the interior of the scale during development, and that chitin pillars form within these invaginations within the cell, but topologically outside it. Furthermore, we found that Calreticulin, a multifunctional protein, is essential for the formation of these trabeculae. This protein was found colocalized with chitin outside the cell membrane, as scales were developing, and its knockout led to loss of chitin pillars, disruption of other scale morphologies, and loss of pigmentation. Our results implicate this multifunctional protein in butterfly wing scale coloration and morphology.

## Introduction

One of the characteristics of butterflies and moths is their colorful and complex wing scales. These scales can contain pigments (Giraldo and Stavenga 2008, Zhang, Martin et al. 2017, Matsuoka and Monteiro 2018, Forman and Thulin 2022) and can be sculpted in elaborate ways to reflect brilliant colors (Ghiradella 1991, Vukusic and Sambles 2003, Nishida, Adachi et al. 2023, Finet, Bei et al. 2024). Each scale is the dried cuticular skeleton of a scale cell and this skeleton contains chitin, an N-acetylglucosamine polymer produced by chitin synthase, an enzyme normally attached to the plasma membrane of cells (Dinwiddie, Null et al. 2014). The scale grows outward during the pupal stage of development. Newly grown-out scales look like baseball bats, but they gradually flatten into wider and thinner paddle-like structures (Dinwiddie, Null et al. 2014, McDougal, Kang et al. 2021, Lloyd, Burg et al. 2024). When scale development is completed, just before adult eclosion, scale cells die leaving behind their cuticle skeleton.

The archetypal skeleton of a butterfly scale is composed of the upper surface, which is mainly assembled from a grid of longitudinal ridges and transverse crossribs, a lower lamina, and pillar-like trabeculae, which connect the upper surface to the lower lamina (Ghiradella and Radigan 1976). In some species, the lumen of the scale is filled with cuticle with more complex and elaborate morphologies. For example, colorful scales of some Papilionidae show a honeycomb lattice between ridges on the upper surface, and some Lycaenidae have perforated multilayer lattices attached to their lower lamina (Ghiradella and Radigan 1976, Ghiradella 1984, Ghiradella 1985, Prum, Quinn and Torres 2006, Seah and Saranathan 2023). What is unique about scales, relative to other types of insect cuticles, is that these cuticular structures are found inside (in the lumen) of dead scale cells. This is mysterious, as cuticle is usually deposited to the outside of the plasma membrane of cells (Binnington and Retnakaran 1991, Merzendorfer and Zimoch 2003, Merzendorfer 2006, Klowden 2013, Sobala, Wang and Adler 2015, Zhu, Merzendorfer et al. 2016, Liu, Cooper et al. 2019) and is here the focus of our investigation.

Early electron microscopy observations of *Mitoura grynea* butterfly pupal wings indicated that cellular membranes were closely associated with cuticle deposition inside the lumen of scale cells with a complex gyroid-shaped cuticular skeleton (Ghiradella 1989). Transmission electron microscopy (TEM) images showed dense crystallite-like structures, surrounded by two membranes, interpreted as the plasma membrane (PM) and the smooth endoplasmic reticulum (SER) membrane (Ghiradella 1989). A chitin deposition model was then proposed where SER budding off the endoplasmic reticulum (ER) helped invaginate and scaffold the PM into the cell lumen, for it to secrete the gyroid cuticular skeleton (Ghiradella 1989). This model, however, has not been validated as no PM nor SER markers were used to distinguish these cellular membranes or to identify the position of these membranes relative to cuticle deposition.

Different visualization techniques have been applied to study the morphogenesis of scales. Traditionally, pupal wings are dissected and fixed at different developmental time points, and visualized with electron microscopy (EM) and confocal imaging (Ghiradella 1989, Ghiradella 1994, Dinwiddie, Null et al. 2014, Day, Hanly et al. 2019, Seah and Saranathan 2023, Lloyd, Burg et al. 2024). More recently, real-time in vivo imaging technology was adopted to further explore scale development (Iwata, Ohno and Otaki 2014, Hirata and Otaki 2019, McDougal, Kang et al. 2021, Nakazato and Otaki 2023, Nakazato and Otaki 2023). However, the limited resolution of conventional confocal fluorescence microscopes and the accumulation of dense cuticle, as well as the lack of adequate stains in EM imaging, have hampered a detailed understanding of scale development. To date, thus, the process of how a stereotypical scale forms, how cuticle gets to be deposited on the inside of a scale cell to form simple trabeculae, and which proteins participate in this process, are still unknown.

To understand how cuticle is being deposited inside a scale cell, we performed immunofluorescence on ultrathin sections of *Bicyclus anynana* butterfly wings, whose scales have an archetypal structure. Simple cuticular pillars, instead of complex gyroids, transverse the lumen of the scale from top to bottom. We imaged these scales with a super-resolution microscope to colocalize plasma membrane and chitin, the main component of insect cuticle. By using immunogold labeling, we found Calreticulin, an ER resident multifunctional protein, located outside the scale cell membrane, and within the cuticular matrix. A CRISPR knockout revealed that Calreticulin is required for the formation of trabeculae, crossribs and other scale nanostructures. Our work discovered an essential role of Calreticulin in scale cuticular structure development.

## Results

### Calreticulin colocalizes with chitin in *B. anynana* wing scales

Complex membrane-cuticle lattice structures are forming around 70% of pupal development (Ghiradella 1989). Thus, we selected 120-hour and 136-hour after pupa formation (APF) (77 and 88% development, respectively) to investigate trabeculae formation in *B. anynana* wing scales. An antibody against the classic ER marker Calreticulin (Moretti, Roy et al. 2017, de la Roche, Hamilton et al. 2018, Wan, Wang et al. 2020, Wang, Christenson and Kinsey 2022) was then used in an immunogold labeling reaction (Fig. 1A-A’’’). Calreticulin showed clear colocalization with the cuticular matrix of wing scales, as the gold particles were found inside ridges, crossribs, trabeculae, and upper and lower laminae in both 120-hour (Fig. 1B-B’’’) and 136-hour APF wing scales (Fig. 1C-C’’’). This suggests that Calreticulin is found outside the ER and outside the scale cell membrane. We were unable to localize the ER further in the scales because other commercial ER markers, such as antibodies targeting Calnexin, were all synthesized using human immunogens with low sequence identity to *B. anynana* orthologous proteins.

**Figure 1.**
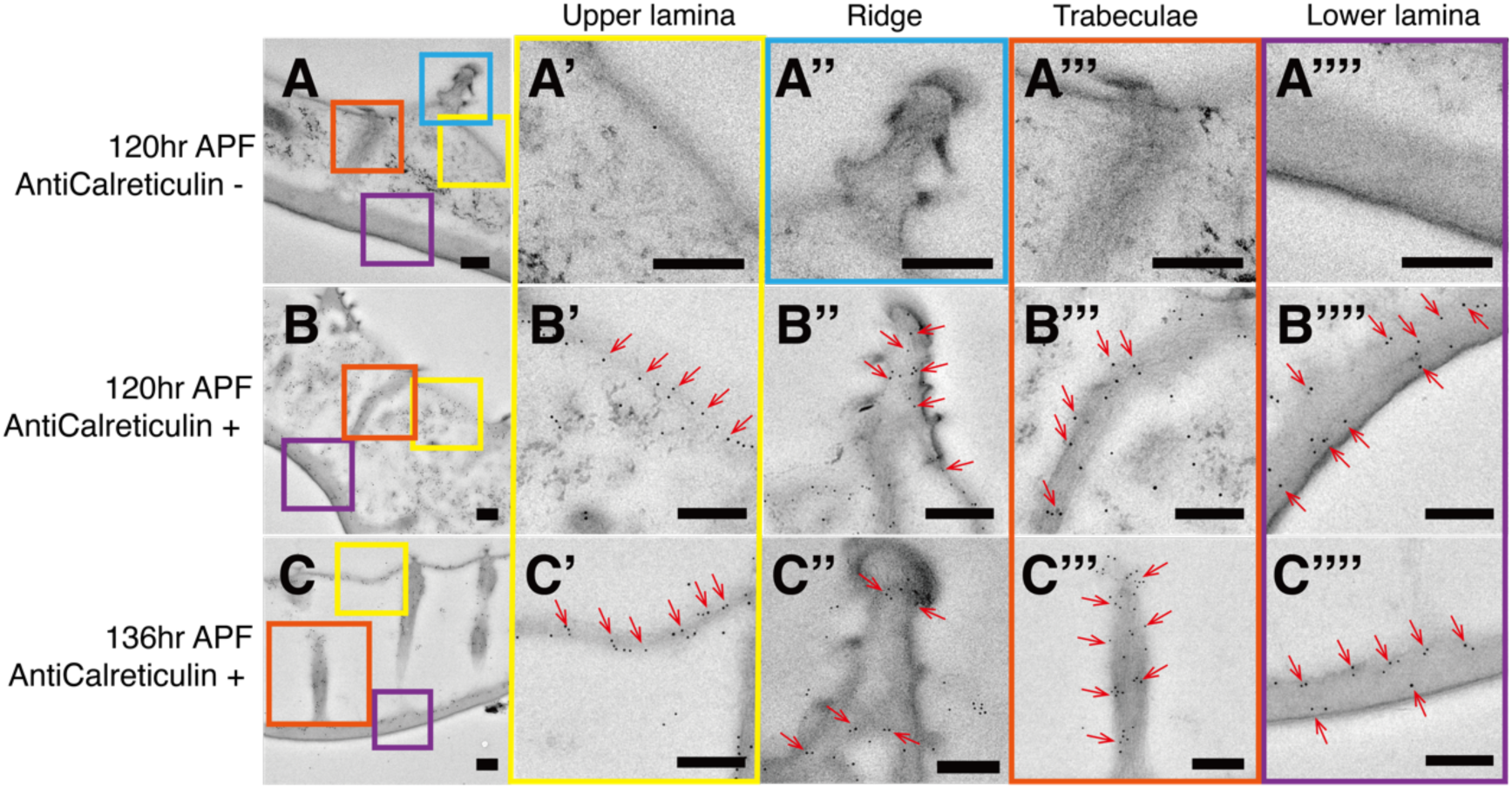
Calreticulin is localized in the cuticular matrix, both on the surface and inside scale cells of *B. anynana*. (A-A’’’’) Immunogold negative control staining on a 120-hour APF pupal wing section. Immunogold staining of Calreticulin on a 120-hour APF wing section (B-B’’’’) and a 136-hour APF wing section (C-C’’’’). Colloidal gold size: 5nm. Enlarged view of different ultra-structures: upper lamina (A’-C’); ridge (A’’-C’’); trabecula (A’’’-C’’’); lower lamina (A’’’’-C’’’’). Red arrows indicate the immunogold particles. Scale bars: 200 nm.

### Chitin and Calreticulin are located topologically outside the scale cell

To colocalize the PM and the cuticular matrix we used immunofluorescence on ultrathin resin-embedded wing sections and two labeled antibodies. We used a house-made antibody against *B. anynana* Armadillo (Banerjee, Murugesan et al. 2023) to visualize the PM, and Wheat Germ Agglutinin (WGA) conjugated with a fluorescent marker to visualize the cuticular matrix. WGA is commonly used to stain chitin (Dinwiddie, Null et al. 2014), a core component of the cuticular matrix, and Armadillo has been used as a marker for the apical cell membrane in an insect chitin deposition study (De Giorgio, Giannios et al. 2023). The tissue sections were first examined under a conventional confocal microscope, but the resolution of these images did not allow us to distinguish the localization of chitin and cell membranes. However, Armadillo did show a positive signal in the scale cells, compared with the negative control preparation, where the antibody against Armadillo was left out (Fig. 2A, B). We then switched to STED super-resolution microscopy (Hell and Wichmann 1994) to improve image resolution. Despite some background noise, the visualization of chitin was clear: WGA stained the typical scale cuticular nanostructures, such as ridges and upper and lower laminae (Fig. 2C-C’’’). Armadillo signals were visible throughout the interior of the scale cross sections (Fig. 2C-C’’’), and sometimes right next to (and around) chitin that was being deposited on the inside (but topologically on the outside) of a scale cell (Fig. 2C’, pink and blue arrowheads). Consistent with this labeling, a structure resembling PM was identified in 131-hour APF wing sections stained with lead citrate under cross-sectional TEM. This membrane surrounded the trabeculae on both sides (Fig. 2D).

**Figure 2.**
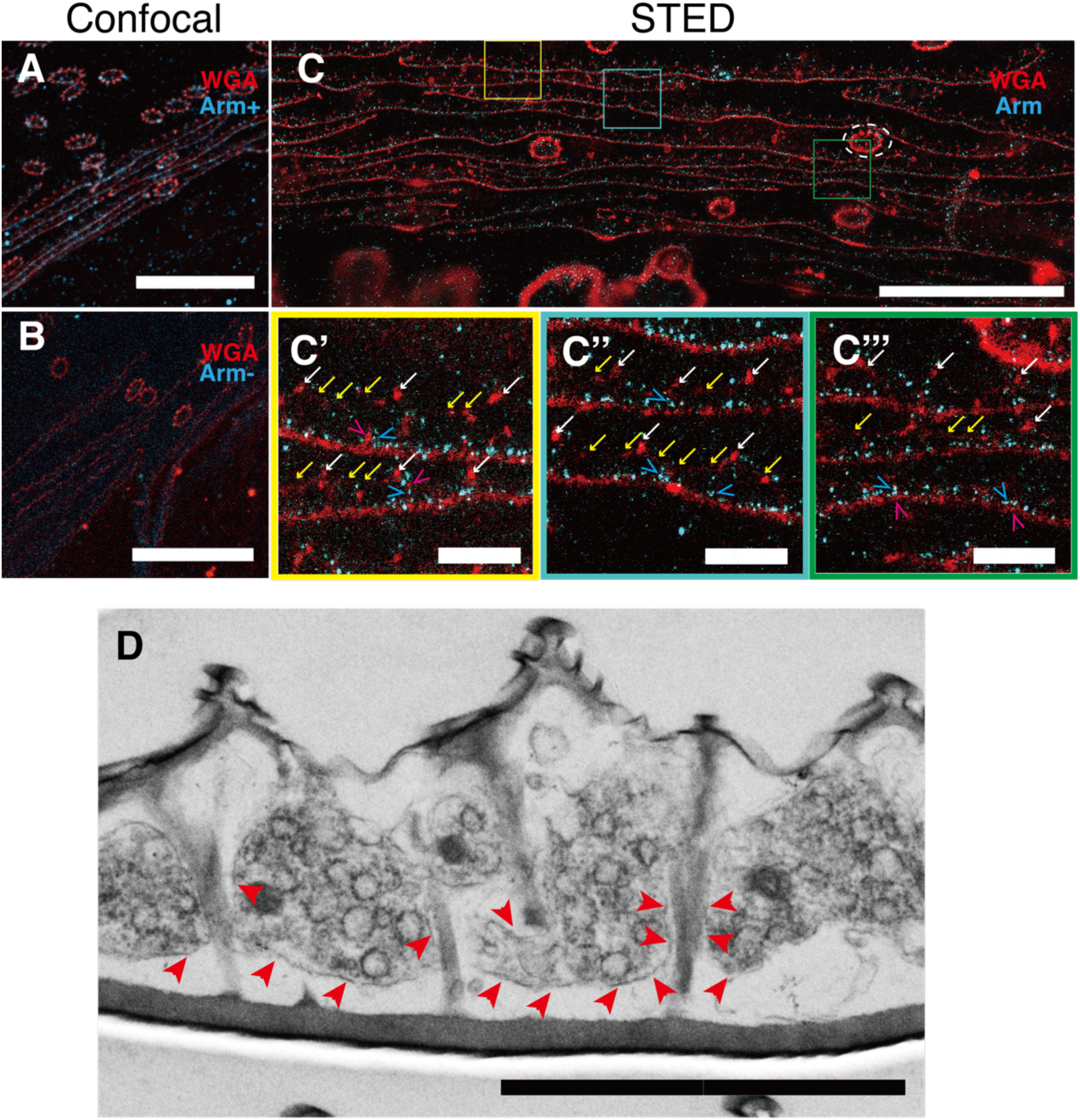
Chitin fingers (WGA staining) are found on the interior of brown scale cells often surrounded by plasma membrane (Arm staining). (A) Confocal image of Armadillo and WGA immunostainings on a *B. anynana* 136-hour APF wing section. (B) A negative control lacking the primary antibody against Armadillo. (C-C’’’) STED super-resolution images of Armadillo and WGA. (C’-C’’’) Enlarged-view images of the regions highlighted in (C). White arrows: ridges. Yellow arrows: upper lamina. Blue arrowheads: Armadillo functions as a plasma membrane marker that surrounds or is tightly associated with WGA-labeled chitin (Pink arrowheads) in vertical trabecula-like structures that are contiguous with the chitin on the outside of the cell (C’), and present across the lumen of scales (C’’), and along the lower lamina (C’’’). White dash circle in (C) shows cross-section of a scale stem. (D) Cross-sectional TEM image of a 131-hour APF wing section. Red arrowheads indicate a plasma membrane-like structure. Scale bars: (A-C): 20 μm. (C’-C’’’, D): 2 μm.

### Knockout of *calreticulin* impacts pigmentation and scale morphology of brown and black scales

As Calreticulin was found in the cuticular matrix we decided to test its function in scale development. We designed a single guide RNA (sgRNA) to target the N-terminus of the transcript using CRISPR-Cas9 (Cal-N, Fig. 3A, Fig. S1). Considering that this protein is indispensable for embryonic development (Michalak, Groenendyk et al. 2009), another sgRNA was designed to disrupt the C-terminal domain (Cal-C, Fig. 3A, Fig. S1), as frame-shifting mutations in this domain lead to the survival of human cells (Klampfl, Gisslinger et al. 2013, Nangalia, Massie et al. 2013, Pietra, Rumi et al. 2016).

**Figure 3.**
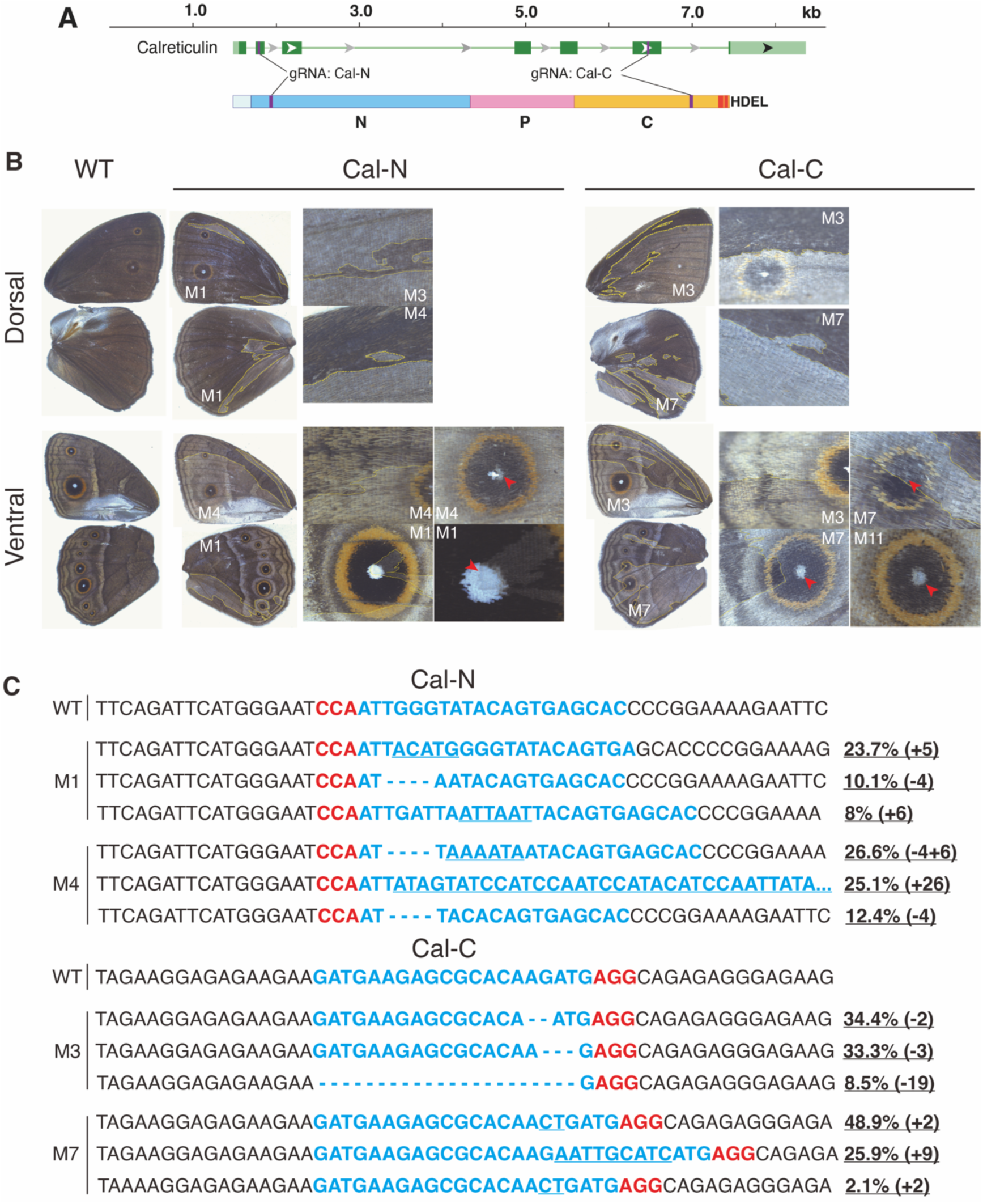
*calreticulin* gene structure and mosaic crispants show loss of melanin pigmentation and partial loss of white scales in the eyespot centers of *B. anynana*. (A) Schematic of *B. anynana calreticulin* gene and encoded protein. N, P and C indicate the N-, P-, and C-domains of Calreticulin. HDEL indicates the motif which prevents secretion from ER. Guide RNA targeting sites are shown in purple. Red bars indicate motifs which are highly conserved across Lepidoptera. (B) Left column: a wildtype (WT) male *B. anynana* dorsal and ventral side views. Middle column: Cal-N crispants. Higher magnification images show color change of scales in mutant wing areas. Right column: Cal-C crispants. Both Cal-N and Cal-C crispants show loss of eyespot center white scales (red arrowheads). (C) The top three most abundant indel alleles in the wings of two representative mutants for each guide RNA. Blue indicates the regions targeted by the guide RNA. Red indicates the protospacer adjacent motifs (PAM). Hyphens indicate deletions. Underlined bases indicate insertions.

Both guides led to similar mutant phenotypes but had different success rates. As expected, embryos injected with Cal-N targeting the N-terminus were mostly inviable: among the 251 larvae that hatched (hatching rate: 43.4%), only 10 larvae survived till adulthood (eclosion rate: 4%). Of those adults, five (50%) showed mosaic phenotypes (Fig. 3B, Fig. S2). Survival was higher with Cal-C targeting the C-terminal domain: among the 162 hatched larvae (hatching rate: 45.9%), 36 eclosed into butterflies (eclosion rate: 22%), and 23 showed mosaic phenotypes (64%) (Fig. 3B, Fig. S2). Both Cal-N and Cal-C crispants exhibited paler scales on both dorsal and ventral sides of wings, regardless of gender (Fig. 3B), with less defined crossribs, whereas most of the white scales in the eyespot centers were missing (Fig. 3B). Next-generation sequencing (NGS) of the affected wing tissue from some of these mosaic mutants confirmed that *calreticulin* was targeted at the two expected gene locations (Fig. 3C, Supplementary data (Cal-C M1.xlsx - Cal-N M4S.xlsx)). With one exception, the frequency of indels causing frameshifts was larger than 50% (ranging from 22-93%; Table 1), suggesting that these mutant wings resulted from either homozygous or hemizygous mutations which contain two different mutated alleles (Table 1, Supplementary data (Cal-C M1.xlsx - Cal-N M4S.xlsx)).

**Table 1.** Summary of mutant allele reads at the *calreticulin* loci targeted by the two guides. Cal-N M4S indicates the genomic DNA sample extracted from the silver area of a Cal-N M4 forewing. Sequencing details can be found in Supplementary data (Cal-C M1.xlsx - Cal-N M4S.xlsx).

| samples | Total reads | Non-frameshifting reads | Frameshifting reads | % reads causing non-frameshifts | % reads causing frameshifts |
| --- | --- | --- | --- | --- | --- |
| Cal-N M1 | 8043 | 2612 | 5431 | 32.5% | 67.5% |
| Cal-N M3 | 9341 | 7294 | 2047 | 78.1% | 21.9% |
| Cal-N M4 | 8812 | 1306 | 7506 | 14.8% | 85.2% |
| Cal-N M4S | 9555 | 630 | 8925 | 6.6% | 93.4% |
| Cal-C M1 | 8209 | 3105 | 5104 | 37.8% | 62.2% |
| Cal-C M3 | 8443 | 4097 | 4346 | 48.5% | 51.5% |
| Cal-C M7 | 11289 | 3969 | 7320 | 35.2% | 64.8% |
| Cal-C M10 | 6184 | 990 | 5194 | 16.0% | 84.0% |
| Cal-C M11 | 9579 | 1797 | 7782 | 18.8% | 81.2% |
| Cal-C M12 | 8741 | 2021 | 6720 | 23.1% | 76.9% |
| Cal-C M13 | 9444 | 4706 | 4738 | 49.8% | 50.2% |
| Cal-C M14 | 8026 | 1535 | 6491 | 19.1% | 80.9% |
| Cal-C M22 | 7664 | 1664 | 6000 | 21.7% | 78.3% |

To quantify the color changes observed in the two types of crispants, we sampled and examined WT and knockout (mKO) scales from mutant individual #4 obtained from Cal-N CRISPR (Cal-N M4) and individual #7 obtained from Cal-C CRISPR (Cal-C M7). We first carried out absorbance measurements in both cover and ground scales from three different colored regions of the wing: dorsal brown scales, ventral brown scales, and ventral eyespot black scales. All mKO scales showed a decrease in absorbance relative to WT scales sampled from the same region (Fig. S3A’’-F’’, Fig. S4A’’-F’’). However, the absorbance spectrum curves did not change in shape, indicating that the mKO scales contained a similar pigment composition but in lower concentration. Optical microscopy images confirmed these findings (Fig. S3A-F, A’-F’, Fig. S4A-F, A’-F’). Mutations in the C- or N-terminal end of the gene produced the same phenotypes (Fig. S3, Fig. S4A-F’’) suggesting that in both cases, Calreticulin’s function in scale coloration was impaired similarly.

To investigate whether mKO scales were impaired in their morphology we used SEM to examine the scale’s surface, top-down TEM to view the scale’s internal structure from the top, and FIB-SEM to obtain a cross-section of the scales. Strikingly, the crossrib morphology which has a typical grid pattern in WT scales (Fig. 4A-A’’) was significantly altered in mKO scales, forming a beaded-bracelet shape between ridges (Fig. 4B-B’’, Fig. S4G, H), or becoming more irregular (Fig. 4B, Fig. S4G, H). Also, unlike WT scales, where open windows are found between the ridges and cross-ridges (Fig. 4A, A’’), most of the mutant scales had their windows closed with a thin cuticular film (Fig. 4B, B’’, Fig. S4G, H). These changes were observed in both Cal-N M4 and Cal-C M7 mutants. Trabeculae in the mKO scales were also shorter or missing, resulting in the upper surface structures (ridges, crossribs, and window-covering film) becoming attached to the lower lamina in places (Fig. 4B’’).

**Figure 4.**
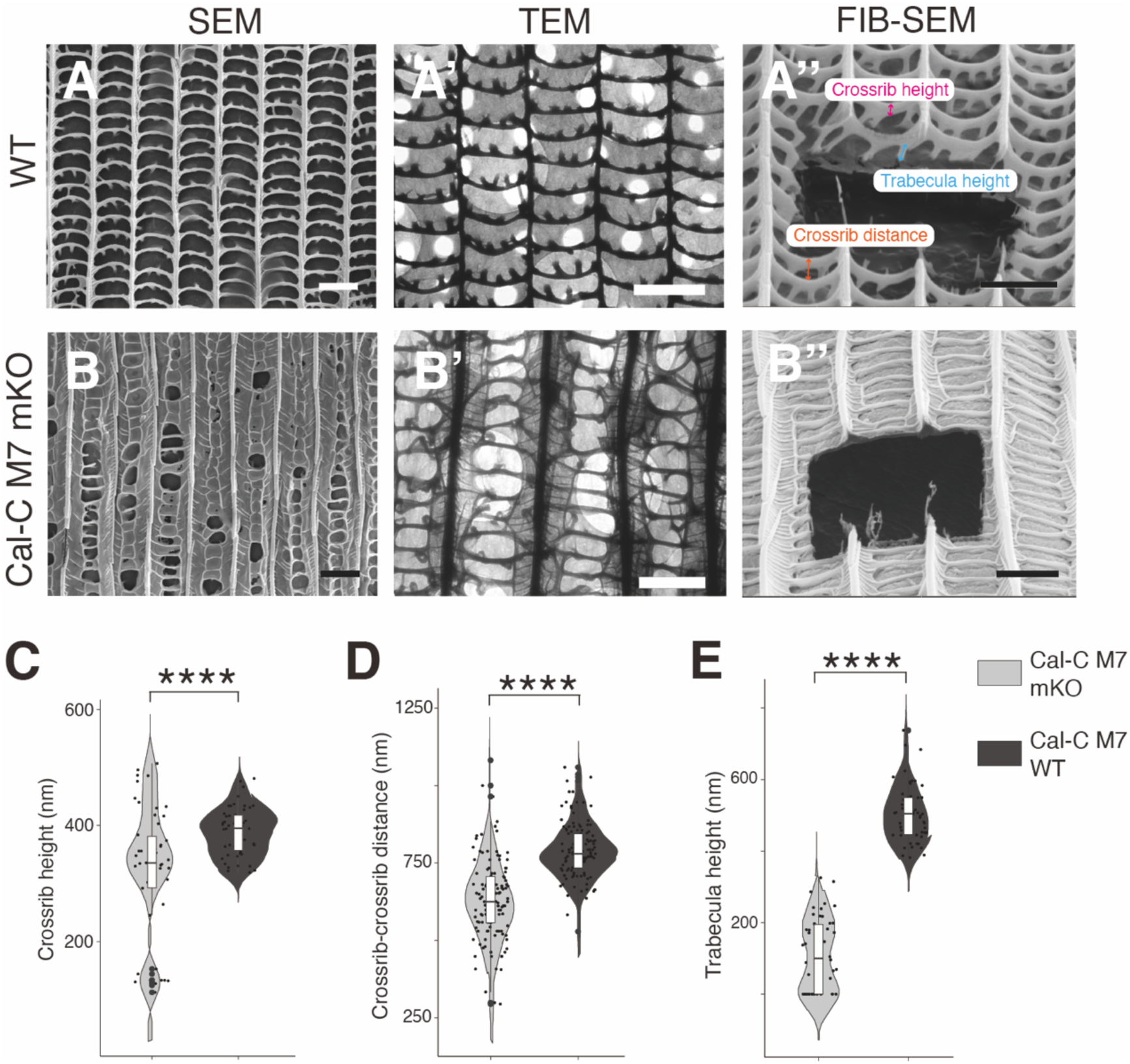
*calreticulin* knockout causes morphological changes in brown cover scales. (A, B) SEM images of the WT and Cal-C M7 mKO dorsal brown scales. (A’, B’) Top-down TEM images. (A’’, B’’) FIB-SEM images of a cross-section of the scales. Scale bars: 2 μm. (C-E) Violin plots of Cal-C M7 cover scale crossrib and trabecula heights, and adjacent crossrib distance. Crossrib height and trabecula height: n = 5, measurements = 50, crossrib-crossrib distance: n = 5, measurements = 125. ****: p < 0.0001. The central line in the violin plot indicates the median of the distribution, while the top and bottom of the box represent the third and first quartiles of the data, respectively. The whiskers show up to 1.5 times the inter-quartile range.

To quantify the structural changes in both cover and ground scales, we measured the height of ridges, crossribs, and trabeculae, the thickness of the lower lamina, and the distance between adjacent crossribs and ridges (Fig. 4A’’, Fig. S5). We found that crossrib height (Fig. 4C) and trabecula height (Fig. 4E) were significantly shorter in the mKO scales, as was the distance between adjacent crossribs (Fig. 4D), whereas the other scale parameters were largely unchanged (Fig. S5).

### Orange, white, and silver scales were not visibly affected upon *calreticulin* knockout but were altered in their morphology

We subsequently investigated changes in the color and nanomorphology of three other scale types. Orange scales are found in the outer rings of the eyespots, silver scales are found in male androconial patches around the scent glands (Prakash, Finet et al. 2022), and white scales are found in the center of the eyespots, most of which disappeared altogether in *calreticulin* mutants (Fig. 3B).

The orange and silver scales did not appear to have changed in color after *calreticulin* disruption (Fig. S6D-G’’, Fig. 5E-E’’, Fig. S2). However, SEM and TEM images showed that mutant orange scales changed in morphology similarly to brown scales (Fig. S6A-C’), whereas silver scales showed no visible morphological or color changes in any of the mutants obtained in this study (Fig. 5E-E’’, Fig. S2). Genetic data showed that silver scales in one of the Cal-N M4 forewings were genetically altered for *calreticulin* with a frame-shifting indel frequency of 93.4% (Table 1) yet showed no significant changes relative to WT silver scales in overall appearance (Fig. 5A, B), morphology (Fig. 5C-C’’, D-D’’), or color (Fig. 5E-E’’). FIB-SEM measurement analysis showed, however, that in Cal-N M4 mKO silver scales, the upper and lower laminae became thicker, and the ridge-ridge distance became wider (Fig. 5C’’, D’’, G-I). WT silver scales display unevenly distributed microribs on a closed upper lamina (Fig. 5C), instead of crossribs and open windows, as seen in all other colorful scales (Fig. 4A, Fig. S6A). They also have web-like trabeculae connecting the upper and lower surface instead of pillar-like trabeculae (Fig. 5C’, C’’). The key parameter that influences the reflectance of silver scales is the air layer thickness (Prakash, Finet et al. 2022) and this was not altered in Calreticulin knockouts (Fig. 5F), which explains why the appearance of these scales remained the same.

**Figure 5.**
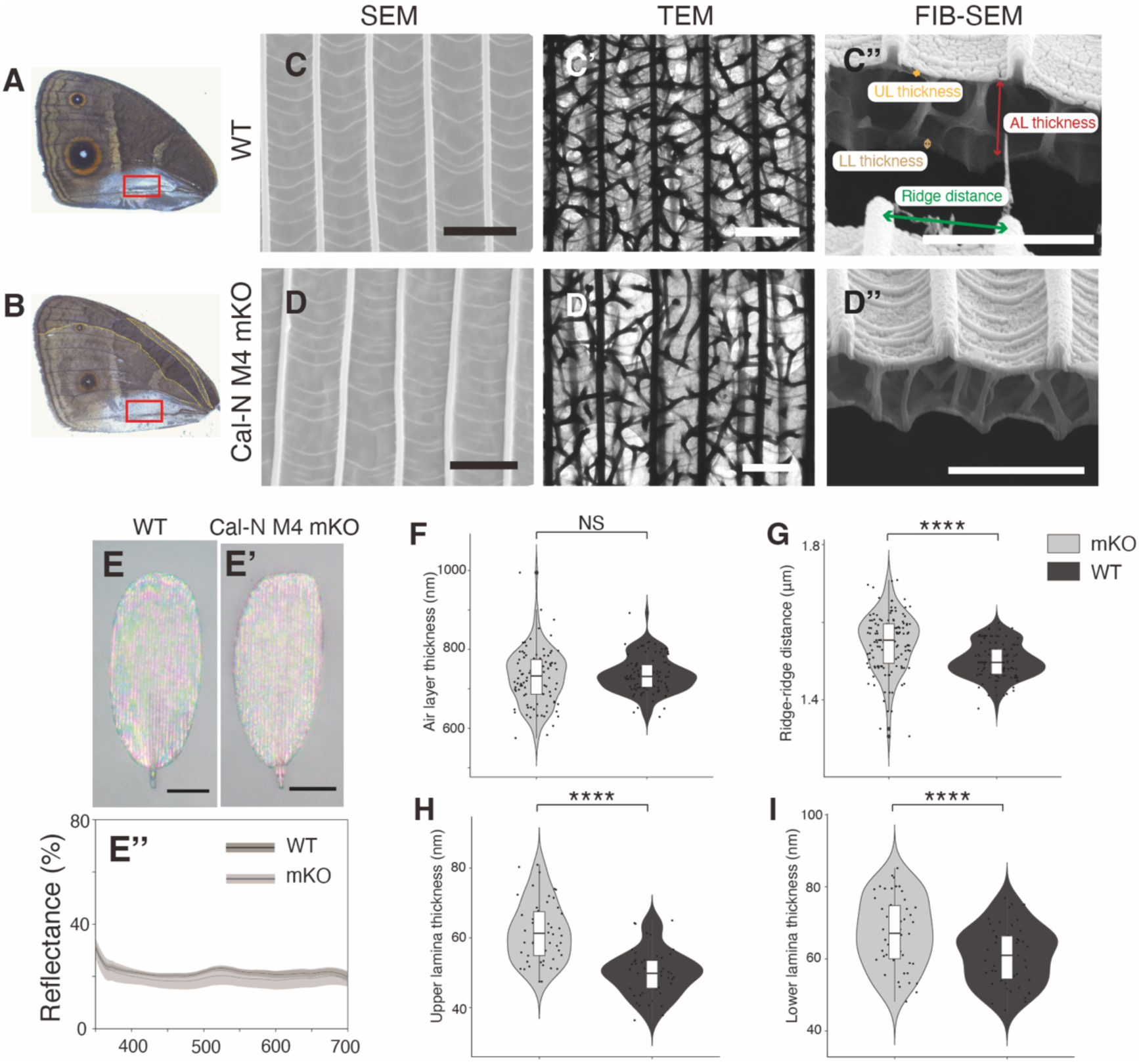
Calreticulin knockouts cause slight alterations of silver scales, but reflectance spectra were not affected. SEM, TEM, FIB-SEM and optical microscopy images of WT (C-C’’, E) and Cal-N M4 mKO silver scales (D-D’’, E’) taken from the red-rectangle demarcated areas in A and B, respectively. (E’’) Reflectance spectra of the mKO silver scales exhibits similar patterns as WT. Scale bars: (C-D’’) 2 μm, (E, E’) 20 μm. (F-I) Violin plots of Cal-N M4 mKO and WT silver scales air layer (AL) thickness, upper and lower lamina thicknesses, and adjacent ridge-ridge distance. air layer thickness: n = 5, measurements = 100; upper and lower lamina thickness: n = 5, measurements = 50; adjacent ridge distance: n = 5, measurements = 125. ****: p < 0.0001, NS: not significant. The central line in the violin plot indicates the median of the distribution, while the top and bottom of the box represent the third and first quartiles of the data, respectively. The whiskers show up to 1.5 times the inter-quartile range.

To investigate why white scales were affected by *calreticulin* mutations we used SEM and FIB-SEM to image their morphology in WT individuals (Fig. 6A) and in Cal-N M4 by picking those residual white scales from the forewing eyespot centers (Fig. 3B, Fig. 6B). This type of scale has a grid-like upper surface nanostructure, similar to the colorful scales, but with a few more partially closed windows, and has long, irregular trabeculae projecting downwards from the crossribs (Fig. 6C, C’). Calreticulin knockouts led to disordered crossribs, pointing in all directions, making the distance between adjacent crossribs impossible to measure (Fig. 6D). In most areas, the scale lumen completely disappeared (Fig. 6D’), and the trabecula height was close to zero (Fig. 6F).

**Figure 6.**
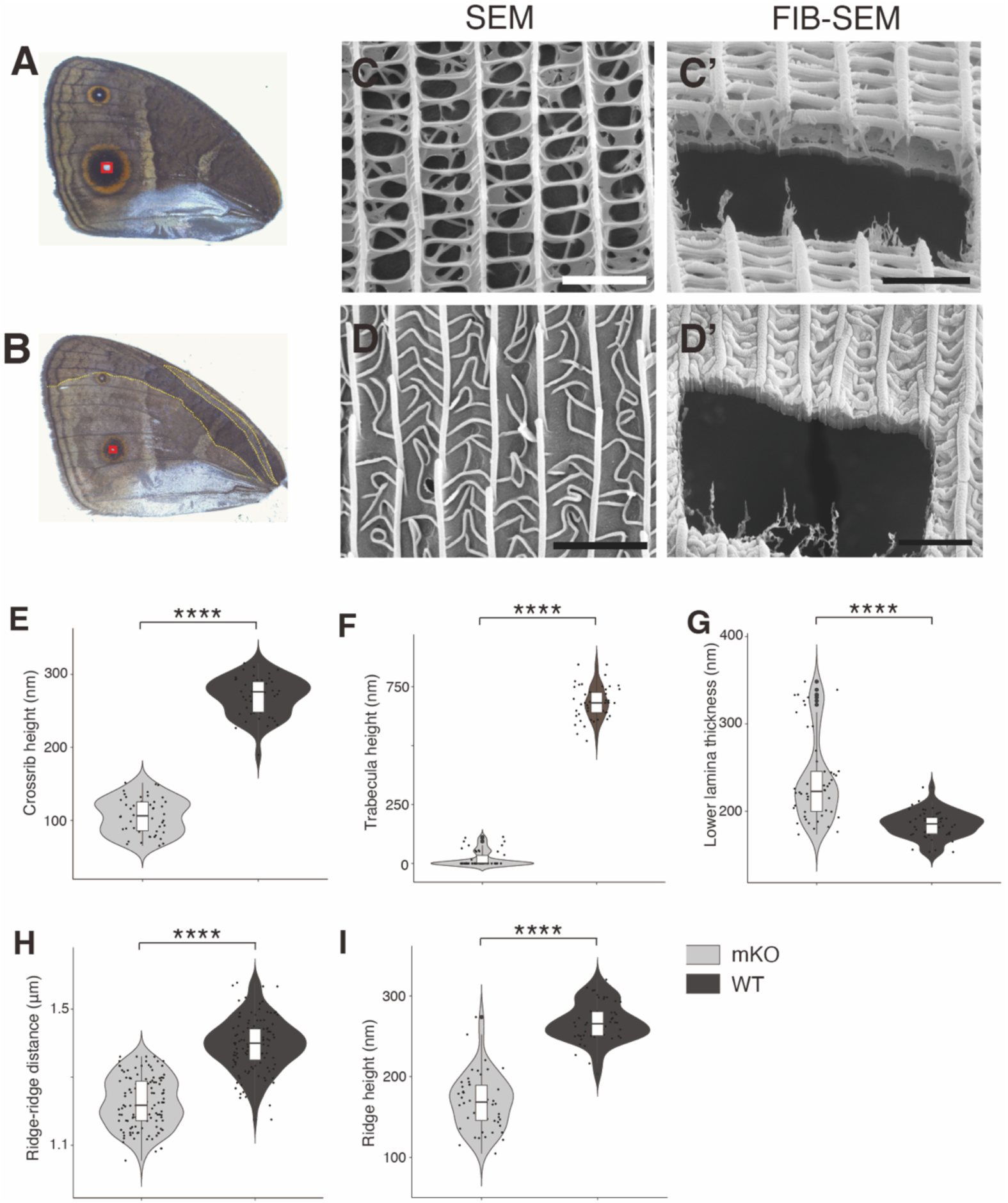
WT eyespot-center white scale morphology was significantly distorted upon *calreticulin* knockout. SEM and FIB-SEM images of WT (C, C’) and Cal-N M4 mKO white scales (D, D’) were taken from the red-rectangle demarcated areas in A and B, respectively. Scale bars: 2 μm. (E-I) Violin plots of Cal-N M4 mKO and WT white scales crossrib and trabecula heights, lower lamina thickness, distance between adjacent ridges, and ridge height. Crossrib and trabecula heights, lower lamina thickness, and ridge height: n = 5, measurements = 50; adjacent ridge distance: n = 5, measurements = 125. ****: p < 0.0001. The central line in the violin plot indicates the median of the distribution, while the top and bottom of the box represent the third and first quartiles of the data, respectively. The whiskers show up to 1.5 times the inter-quartile range.

All these data indicate that *calreticulin* primarily impacts the development of scales that have well-defined crossribs, and associated trabeculae, such as white, orange, brown, and black scales, but not silver scales.

### Lepidopteran Calreticulin

Guide RNA Cal-C only influenced a short sequence at the C-terminal end (Fig. S1). However, the phenotype it produced was the same as Cal-N crispants. To explain this result, we sought to identify motifs that could function in scale cuticular development at the C-terminal end of *B. anynana*. Only a single isoform of *calreticulin* was found in the *B. anynana* genome and transcriptome (https://www.ncbi.nlm.nih.gov/gene/112045067). To compare the amino acid sequence of this protein to other orthologs, we aligned Calreticulin sequences across four insect orders (Lepidoptera, Diptera, Coleoptera, and Hymenoptera). Whole sequence alignment of Calreticulin indicates that the C-terminal ends vary among species (Fig. S7A, Fig. S8A, Fig. S9A - Fig. S12A) and different orders have different C-terminal end patterns (Fig. S7B, Fig. S9B - Fig. S12B). Two conserved and unique motifs were found at the C-terminal end of Lepidopteran sequences (Fig. S7B, Fig. S9). Alignment results as FASTA Alignment output files can be found in the Supplementary data-FASTA alignment.zip.

## Discussion

In this study, we found evidence that the cuticular pillars (trabeculae) that transverse the lumen of dead scale cells, are likely formed by cuticle being deposited to the outside of finger-like invaginations of the plasma membrane in developing scale cells. Our STED and TEM data support the idea that these pillars are formed outside the scale cell, topologically, while clearly transversing it from one side to the other. We also discovered that the classic ER-resident protein Calreticulin cannot be used as an ER membrane marker at the stages of scale development examined here (77-88% of pupal development), as this protein was found embedded in the cuticle of *B. anynana* wing scales, both in the upper and lower laminae, and inside the trabeculae that transversed the scales (Fig. 1).

Mutations targeting *calreticulin* significantly altered the morphology of all the scales with well-defined crossribs (brown, black, orange and white scales) (Fig. 4, Fig. S4G, H, Fig. S6A-C’, Fig. 6C-D’) and made brown and black scales (but not orange scales) lose pigmentation (Fig. S3, Fig. S4A-F’’, Fig. S6D-G’’). In general, mutations in *calreticulin* affected the formation of crossribs and their orientation, and inhibited the formation of trabeculae below the crossribs, making them shorter or missing (Fig. 4, Fig. S5). White scales were most severely affected by these mutations. They had severely disordered crossribs (Fig. 6D) and an absent scale lumen (Fig. 6D’) with no trabeculae (Fig. 6F). It is likely that white scales formed but were lost upon adult emergence, perhaps due to the large morphological alterations (Fig. 3). In contrast, silver scales, which do not produce crossribs, were only slightly affected in their morphology, and their development does not seem to rely on Calreticulin.

The specific effect of Calreticulin on the development of crossribs and trabeculae could perhaps explain why only brown or black scales, showed alteration in pigmentation, but not orange scales. Orange scales contain primarily ommochrome pigments (How, Banerjee and Monteiro 2023, Banerjee, Finet et al. 2024), whereas brown scales contain melanins (Matsuoka and Monteiro 2018). If ommochromes are preferentially deposited along longitudinal ribs, and melanins preferentially deposited in crossribs and trabeculae (Banerjee, Finet et al. 2024) then, preventing these latter structures from forming would prevent melanins from being deposited in the mass of the scale, leading to lighter scales. An additional explanation for the pigmentation effects is that Calreticulin mutations are known to lead to misfolding of an enzyme in the melanin pathway, tyrosinase, which would affect melanin production (Petrescu, Petrescu et al. 1997, Liedy 2005).

### Trabecula formation

Here we propose a model of trabeculae formation based on our experimental observations. We noted that in *B. anynana* the lower lamina and ridges form first, followed by the crossribs and the pillar-like trabeculae, a pattern previously noted in other species (Ghiradella 1989, Dinwiddie, Null et al. 2014). When crossribs first formed they were thin, but later became thicker. In *calreticulin* mutants, the crossribs remain thin, and it is possible that the shorter or missing trabeculae in these mutants result from failed growth of trabeculae below these thin crossribs. In our model we propose that at the late pupal developmental stage, with the involvement of Calreticulin, crossribs form in between the ridges. Meanwhile, at points along these crossribs, fingers of plasma membrane invaginate towards the interior lumen of the cell, fuse with the membrane on the other side of the scale and form an extracellular channel into which cuticle is deposited to form trabeculae (Fig. 7B, C). We name these points “trabecula starting points” (Fig. 7B). The scale cell then dies leaving its cuticular skeleton behind where the cuticle transverses the scale (Fig. 7D). When *calreticulin* is mutated, crossrib formation is disrupted, impairing trabecula starting points located at crossribs and preventing trabecula from forming (Fig. 7B’, C’). Upon cell death, and without the support of pillar-like trabeculae, the upper surface collapses and is found fused to the lower lamina (Fig. 7D’). The loss of these trabeculae also affects melanin deposition in these structures, perhaps contributing to mKO scales becoming paler.

**Figure 7.**
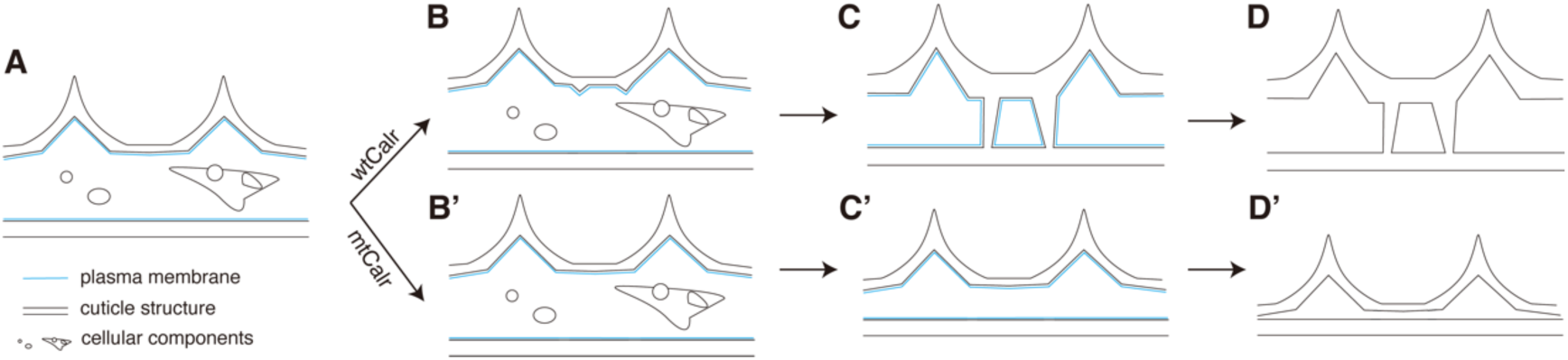
Model for how Calreticulin affects the formation of crossribs and trabeculae. (A) At ∼70% development, ridges and thin crossribs have formed. (B-C) “Trabecula starting points” start to form from further-developed crossribs with the help of WT Calreticulin. (B’-C’) Mutating Calreticulin impairs crossrib formation, which impairs trabecula formation. (D, D’) Cells die after scale development is complete, leaving their cuticular skeleton behind. (D’) Without trabecula support, cell death results in the upper surface structures (ridges and crossribs) becoming attached to the lower lamina.

In silver scales, there are complex trabeculae connecting the two laminae (Fig. 5C’’) but they seem to originate at the lamina and not at the thin microribs. Trabeculae starting points located in the lower or upper laminae (outside the crossribs) may not be affected by Calreticulin dysfunction. Given that *calreticulin* mutations do not affect silver-scale trabeculae morphology, we speculate that Calreticulin is dispensable for the formation of those trabeculae.

### The lectin domain and C-terminal end of Calreticulin are potential functional elements in scale nanostructure development

Calreticulin is a pleiotropic protein that is essential to all eukaryotes. The functions of Calreticulin include but are not limited to calcium buffering, chaperoning glycoproteins, cell adhesion, antigen presentation on the cell surface, initiation of apoptotic-cell removal, and wound healing (Coppolino, Woodside et al. 1997, Ogden, deCathelineau et al. 2001, Gardai, McPhillips et al. 2005, Obeid, Tesniere et al. 2007, Michalak, Groenendyk et al. 2009, Fucikova, Spisek et al. 2021, Sawaya, Vecin et al. 2023). The pleiotropic nature of this protein explains why most injected individuals (96%) in the Cal-N CRISPR died during growth, indicating Calreticulin’s essential function for embryonic survival. However, *calreticulin* mutations did not influence wing cell survival as genotyping results confirmed that most wing cell clones affected were homozygous mutants with frameshifts (Fig. 3C, Table 1, Supplementary data (Cal-C M1.xlsx - Cal-N M4S.xlsx)).

To explain how Calreticulin might impact scale cuticle development we explored its protein sequence in detail. There are three main domains in Calreticulin proteins: N-, P-, and C-domains (Fig. 3A, Fig. S1). N- and P-domains are important for the glycoprotein chaperone function and the C-domain is important for the calcium buffering function (Michalak, Groenendyk et al. 2009, Wang, Groenendyk and Michalak 2012, Fucikova, Spisek et al. 2021). The N-terminus contains a carbohydrate-binding site in a globular lectin domain. Lectins, such as Conconvalina (ConA), WGA, and Soybean agglutinin (SBA), are a group of glycoproteins that can bind glycans or sugars (Ni and Tizard 1996, Kremsreiter, Kroell et al. 2021). Chitin, a polymer of N-acetylglucosamine, is a type of sugar (an amide derivative of glucose). Therefore, the lectin activity of Calreticulin makes it a potential chitin-binding protein, which would explain its presence in the cuticular matrix of developing scales (Fig. 1), and its newly discovered function in scale crossrib and trabecula development.

In this study, however, Cal-C CRISPR mutants showed the same phenotypes as Cal-N mutants, suggesting that another functional element may exist in the C-terminal end (Fig. S1). Calreticulin C-terminal ends vary dramatically across insect orders and lepidopterans exhibit two unique amino acid motifs (Fig. S7B, Fig. S9-Fig. S12). We speculate that these unique motifs might also be involved in scale cuticle development, but future work is required to test this hypothesis. Furthermore, it is worth noting that scales from butterflies in families other than Nymphalidae do not have the typical “ridge-crossrib-trabecula-lower laminae” nanostructures. The interior lumen of these scales can have honeycomb lattices, perforated multilayer lattices, and multiple other cuticular structures (Ghiradella and Radigan 1976, Ghiradella 1984, Ghiradella 1985, Prum, Quinn and Torres 2006, Seah and Saranathan 2023). It might be interesting to explore how Calreticulin functions in the formation of all these nanostructures.

## Supporting information

FASTA alignment.zip

## Acknowledgements

We thank Sim Aik Yong from the Electron Microscopy Unit (EMU) of Yong Loo Lin School of Medicine, NUS, for help with the EM section preparation, post-staining, and TEM operation, Cheng Liyan Sherona from the Department of Chemistry, NUS, for access and help with SEM, the Electron Microscopy Facility (EMF, NUS) for use of FIB-SEM, Abberior GmbH company for use of their super-resolution STEDYCON microscope, E. Chae’ lab and Y. Yun for help with the genotyping NGS, Deepan Balakrishnan for providing a top-down TEM protocol, and Tirtha Das Banerjee for providing the *B. anynana* Armadillo antibody. This research was supported by the National Research Foundation (NRF) Singapore under the Competitive Research Program Program (award NRF-CRP20-2017-0001) and the Ministry of Education, Singapore (award MOE-T2EP30222-0017).

## Author Contributions

H.R. and A.M. conceived and designed the study. H.R. performed all the experiment and collected the spectral measurements and the top-down TEM data. H.R. and C.F. collected the SEM data, and C.F. collected the FIB-SEM data. C.F. did the FIB-SEM measurement and analyzed the data. H.R. wrote the original draft. A.M. reviewed, edited, and completed the manuscript with inputs from all the authors.

## Materials and methods

### Butterfly Husbandry

*B. anynana* butterflies were reared at 27 °C as described in a previous publication (Prakash and Monteiro 2018). To harvest pupal wings at specific developmental time points, the time of pupation was recorded.

### TEM Immunogold Labeling

Butterfly pupal wings were fixed with 4% formaldehyde in 0.1 M PB (0.1 M Phosphate Buffer, pH7.4) for 12-24 hours at 4 ℃, rinsed in 0.1 M PB for 3×10 min and quenched in 0.1 M Glycine in 0.1 M PB. The wings were then incubated in 0.2 M Sucrose in 0.1 M PB for 2×15min at room temperature. Dehydration was performed by incubating wings in two changes of 70% ethanol 30 minutes each, then in LR white resin and 70% ethanol (2:1) mixture for 1 hour, and in three changes of pure LR white resin for 1 hour each. Wings were incubated in the last change of LR white resin overnight at room temperature. Samples were then placed in the bottom of a gelatin capsule, filled with LR white resin to the brim, and polymerized in a 60 ℃ oven for 24-48 hours or longer. Those EM blocks were trimmed and sectioned at 90-100 nm. Sections were rinsed for 2×2 min by placing grids on large droplets of TBS-Tween (0.05 M TBS, 0.05% Tween 20, pH 7.6) and blocked in IHC Ultra Serum Blocking buffer (1% BSA, 3% Normal Serum, 0.1% Fish Gelatin, 0.05% Sodium Azide in 0.05 M TBS, pH 7.6) for 30 minutes. For primary antibody labeling, sections were incubated on droplets of rabbit anti-Calreticulin antibody (ab92516, Abcam) diluted 20 times in Universal Antibody Diluent (ab79995, Abcam), overnight at 4 ℃. This was followed by rinsing on large droplets of TBS-Tween for 6×2 min. For the negative control, the same volume of Universal Antibody Diluent without primary antibody was used. For secondary antibody staining, the sections were incubated on droplets of secondary immunogold-conjugated antibodies (Anti-Rabbit IgG (whole molecule) – This antibody was produced in a goat, and conjugated with 5nm gold nanoparticles (colloidal gold, G7277, Sigma-Aldrich) at 1:40 in 1% serum/TBST + 0.5% polyethylene glycol for 1 hour at room temperature. Sections were post-fixed on droplets of 2.5% glutaraldehyde in 0.1 M PB for 10 minutes and stained with lead citrate for 5 minutes. Images were acquired on a JEOL 1400Flash TEM (JEOL Ltd. Japan).

### Immunofluorescence

To visualize Armadillo proteins in wing scale cross sections, the EM resin blocks generated for immunogold staining were sectioned at 200-300 nm and attached onto glass slides. These sections were rinsed for 2×2 min with TBS-Tween and blocked in IHC Ultra Serum Blocking buffer for 2 hours. For primary antibody labeling, sections were incubated in 1:40 diluted house-made Rat anti-*Bicyclus* Armadillo antibody (Banerjee, Murugesan et al. 2023) in Universal Antibody Diluent (ab79995, abcam) overnight at 4 ℃. This was followed by rinsing on TBS-Tween for 6×2 min. For the negative control, the same volume of Universal Antibody Diluent without primary antibody was used. For secondary antibody staining, 1:50 diluted Goat Anti-Rat IgG H&L (Alexa Fluor® 647) (ab150159, abcam) was applied on the sections for 2 hours. After washing with TBS-Tween for another 6×2 min, the sections were stained with wheat germ agglutinin (WGA) Conjugated with Alexa Fluor™555 (w32464, ThermoFisher) and mounted in Abberior mounting medium. The slides were examined with a Zeiss LSM900 confocal microscope and an Abberior STED microscope (STEDYCON, Abberior) to acquire images.

### CRISPR/Cas9 deletion

The CRISPR knockout method through embryonic injection was described previously (Banerjee and Monteiro 2018). Instead of single guide RNAs (sgRNA), in this study, CRISPR using crRNA-tracrRNA duplex was carried out following the protocol provided by IDT (Integrated DNA Technologies, U.S.A). Cal-N and Cal-C crRNAs were designed and their on-target score assessed by the Custom Alt-R™ CRISPR-Cas9 guide RNA design service, IDT. Both crRNA and tracrRNA were synthesized by IDT. 4 μl Nuclease-Free Duplex Buffer (IDT) was mixed with 0.5 μl 100 μM crRNA and 0.5 μl 100 μM tracrRNA. The mixture was heated at 95 ℃ for 5 min and cooled down to room temperature. 0.5 μl of 10 µg/µL Cas9 protein solution (Alt-R™ S.p. Cas9 Nuclease V3, 1081058, IDT) was diluted in 5 μl Cas9 working concentration buffer (20 mM HEPES; 150 mM KCI, pH 7.5) and mixed with 0.05 μmol crRNA-tracRNA duplex, followed by incubating at 37 ℃ for 5 min to form RNP. The Cas9 RNP complex was then ready for injection. All crRNA and primers used for NGS genotyping were summarized in Table S1.

### Optical imaging and UV-VIS-NIR microspectrophotometry

Light microscope images of individual scales were recorded using the 20X objective lens of a uSight-2000-Ni microspectrophotometer (Technospex Pte. Ltd., Singapore) and a Touptek U3CMOS-05 camera. Scales from either WT or mKO areas were taken, mounted on a glass slide, and viewed using Llumins Toupview microscope camera software. For silver scales, the light source was provided by a Thorlabs Mercury-Xenon Short-Arc Light Source (SLS402, ThorLabs Inc., New Jersey, USA). To obtain better images, only intact scales were chosen and images were taken at different focal distances. Z-stacking was carried out using EDF function (<u>Process>EDF•••)</u> in Toupview. Maximum Contrast with default settings, followed by no Auto Align, was used as the EDF method.

To measure absorbance, scales were mounted in clove oil (C8392, Sigma-Aldrich, refractive index = 1.532), with a similar refractive index to chitin. Matching the reflective indexes minimizes the production of structural colors and allows the quantification of pigmentary-based colors alone. The absorbance spectrum was measured for individual scales using a 20X objective, with 100 ms integration time and 10x averaging. For each scale type, average values were taken from three measurements of each scale, and six to eight individual scales were measured. Spectra with usable range between 400 and 700 nm were shown. Analyses and spectral plots were done in R Studio 2023.03.0 Build 386 with R version 4.2.3 (2023-03-15) (R Core Team (2023) (R Core Team 2013) URL: https://www.R-project.org/.) using the R-package pavo (v 2.7) (Maia, Gruson et al. 2019).

To measure reflectance, an aluminum reflector sheet was used as a reference for 100% reflectance. Individual silver scales from a WT male forewing or the Cal-N M4 forewing were picked and mounted on glass slides and illuminated with the same light source as in the optical imaging of silver scales. Normal-incidence UV-VIS-NIR reflectance spectra with a usable range between 335 and 700 nm were collected using a high NA 100x objective. For each scale, three measurements were taken from three different areas of the scale surface and five to ten scales were measured for each scale type.

### Focused ion beam scanning electron microscopy (FIB-SEM)

FIB-SEM method was described in a previous publication (Prakash, Finet et al. 2022, Banerjee, Finet et al. 2024). Five to ten scales picked from each selected area of the wings were used for each sample. Images were taken with a FEI Versa 3D with the following settings: beam voltage 8kV, beam current 12pA at a 52° tilt. Image acquisition was performed in the same equipment with the following settings: beam voltage 5kV, beam current 13pA.

### Scanning electron microscopy (SEM)

More than 10 scales from each selected area of the wings were picked using insect pins and individually mounted onto carbon tape stuck on a SEM stub. The scales were then sputter coated with platinum using a JFC-1600 auto Fine Coater (JEOL Ltd. Japan). Images were obtained using a JEOL JSM-6701F Field Emission Scanning Electron Microscope (FESEM) (JEOL Ltd. Japan).

### Cross-sectional TEM

Pupal forewings at 131-hour APF were dissected and fixed in 2.5% Glutaraldehyde at 4℃ overnight followed by three washes in PBS (20min for one wash). Regular cross-sectional TEM was performed on stained samples using potassium ferrocyanide in osmium tetroxide for the post-fixation (1.5% Potassium ferrocyanide K₄[Fe(CN)₆] in 1% Osmium Tetroxide) as described in a previous publication (Prakash, Balakrishnan et al. 2023). Images were acquired on a JEOL 1400Flash TEM (JEOL Ltd. Japan).

### Measurements and statistical analysis

Measurements of thicknesses and distances were measured from SEM images using the Line tool implemented in Fiji (Schindelin, Arganda-Carreras et al. 2012). For all parameters, repeated measurements were taken per scale with five scales sampled from one area. Due to the multilevel context of the datasets, we ran linear mixed-effects (LME) models using the R package nlme (Pinheiro, Bates and Team 2023) that allows coefficients to vary with respect to one or more grouping variables. The scale type was treated as the fixed factor, and the scale nested within individual as a random factor. The lack of homogeneity of variances among scale types prompted us to use the varIdent() function in the nlme package. Akaike information criterion (AIC) was used to compare different possible models and determine which one is the best fit for the data. Outcomes of the LME tests and the p-values are given in Supplementary data -Tables S2.xlsx and Tables S3.xlsx. Measurements were plotted using the R package ggplot2 (Gómez-Rubio 2017). For crossrib height, trabecula height, lower lamina thickness and ridge height, for each scale type, 5 individual scales were selected, and 10 individual measurements were taken within a scale. Total measurements = 50. For crossrib distance and ridge distance, for each scale type, 5 individual scales were selected, and 25 individual measurements were taken within a scale. Total measurements = 125.

### Top-down transmission electron microscopy (TEM)

Formvar-coated TEM grids were glow-discharged for 30 seconds, at 5 mA using a Leica EM ACE200. Ten or more individual scales were picked from each selected wing area and placed on the discharged grids. Top-down TEM images were then taken using a JEOL 1400Flash TEM.

### Genotyping adult wings

Genotyping of crispants was carried out using a method described in a previous publication (Tian, Asano et al. 2024). Genomic DNA was extracted from mutant adult wing patches showing both WT and mKO phenotypes except Cal-N M4S, which was extracted from a pure mKO area of a Cal-N M4 forewing covered by silver scales. Next-generation sequencing (NGS) was performed on an Illumina iSeq 100 system, using 2×150bp paired-end (PE) sequencing. Sequence analyses were done using the online web-tool “CRISPR RGEN Tools” (http://www.rgenome.net/). Genotyping results are shown in (Fig. 3C, Table 1, Supplementary data-Cal-C M1.xlsx - Cal-N M4S.xlsx).

### Multiple alignments of insect Calreticulin

The alignment was carried out using COBALT (Constraint-based Multiple Alignment Tool, https://www.ncbi.nlm.nih.gov/tools/cobalt/cobalt.cgi) by uploading the accession ID of Calreticulin proteins. We included 15 representative lepidopteran sequences from Nymphalidae, Papilionidae, Pieridae, Lycaenidae, Pyralidae, Noctuidae, Geometridae, Sphingidae, and Bombycidae; Eight representative dipteran sequences from Culicidae, Tephritidae, Cecidomyiidae, and Drosophilidae; Ten representative coleopteran sequences from Brentidae, Buprestidae, Chrysomelidae, Coccinellidae, Curculionidae, Lampyridae, Nitidulidae, Scarabaeidae, and Tenebrionidae; And eight representative hymenopteran sequences from Ampulicidae, Apoidea, Formicidae, Braconidae, Athaliidae, and Diprionidae. For the alignment of all four orders, all 41 representative Calreticulin sequences were uploaded. The accession ID and species information can be found in Supplementary data-Calreticulin_ACCESSION.xlsx.

