## Supplemental Table 1 and Figures for "Calreticulin is required for cuticle deposition and trabeculae formation inside butterfly wing scale cells"

| | |
|---|---|
| Cal-N (guide RNA) | GTGCTCACTGTATACCCAAT |
| Cal-C (guide RNA) | GATGAAGAGCGCACAAGATG |
| Cal-N-F (sense primer) | CCGGCTTTTGATATTGATTTTTGATG |
| Cal-N-R (antisense primer) | GACAACGTGCCAATTAGAAAGGCGC |
| Cal-C-F (sense primer) | CGGGAACCATCTTCGACAACAT |
| Cal-C-R (antisense primer) | TACAATAGTGCACTCACCACAGGAG |

**Table S1 Guide RNAs and genotyping primers used in this study.**

>XP_023936899.1 calreticulin [Bicyclus anynana]

MKSLVLSLVSLLALYSV*NCEVYYEEKFPDDSWESNWVYSEHPGKEFGKFKLTAGKFYNDPE*

*EDKGLQTSEDARFYALSRKFKPFSNEGKDLVIQFTVKHEQDIDCGGGYVKVFDCNLDQKDM*

*HGESPYEIMFGPDICGPGTKKVHVIFSYKGKNHLIKKDIRCKDDVHTHLYTLVVKPDNKYEVLI*

*DNEKVEDGNLEDDWDFLPPKKIKDPEAKKPE*DWDDRATIPDPDDTKPEDWDKPEHIPDPDA

SKPEDWDDEMDGEWEPPMIDNPDYKGVWSPKQIDNPAYKGAWIHPEIDNPEYTEDKNLYL

RKEICAVGLDLWQVKSGTIFDNILFTDDLEVAKARGEELKKTLEGEKKMKSAQD**EAEREKEK**

**AEKPEEEDDEDLDDEIAPEGESAPVEEHDEL**

**S:Cal-N crispr disrupted start point**

K:Cal-C Crispr disrupted start point

VYYEEKFPDDS…: N domain

DDWDFLPPKKI…: P domain

EICAVGLDLWQ…: C domain

**EAEREKEKAEKPEEEDDEDLDDEIAPEGESAPVEEHDEL: totally different seq from human CALR**

*NCEVYY*: lectin domain

**Figure S1. *B. anynana* Calreticulin protein sequence.** N, P, and C domains as well as the lectin domain were defined by blasting against human Calreticulin. Sites targeted by CRISPR are also highlighted.

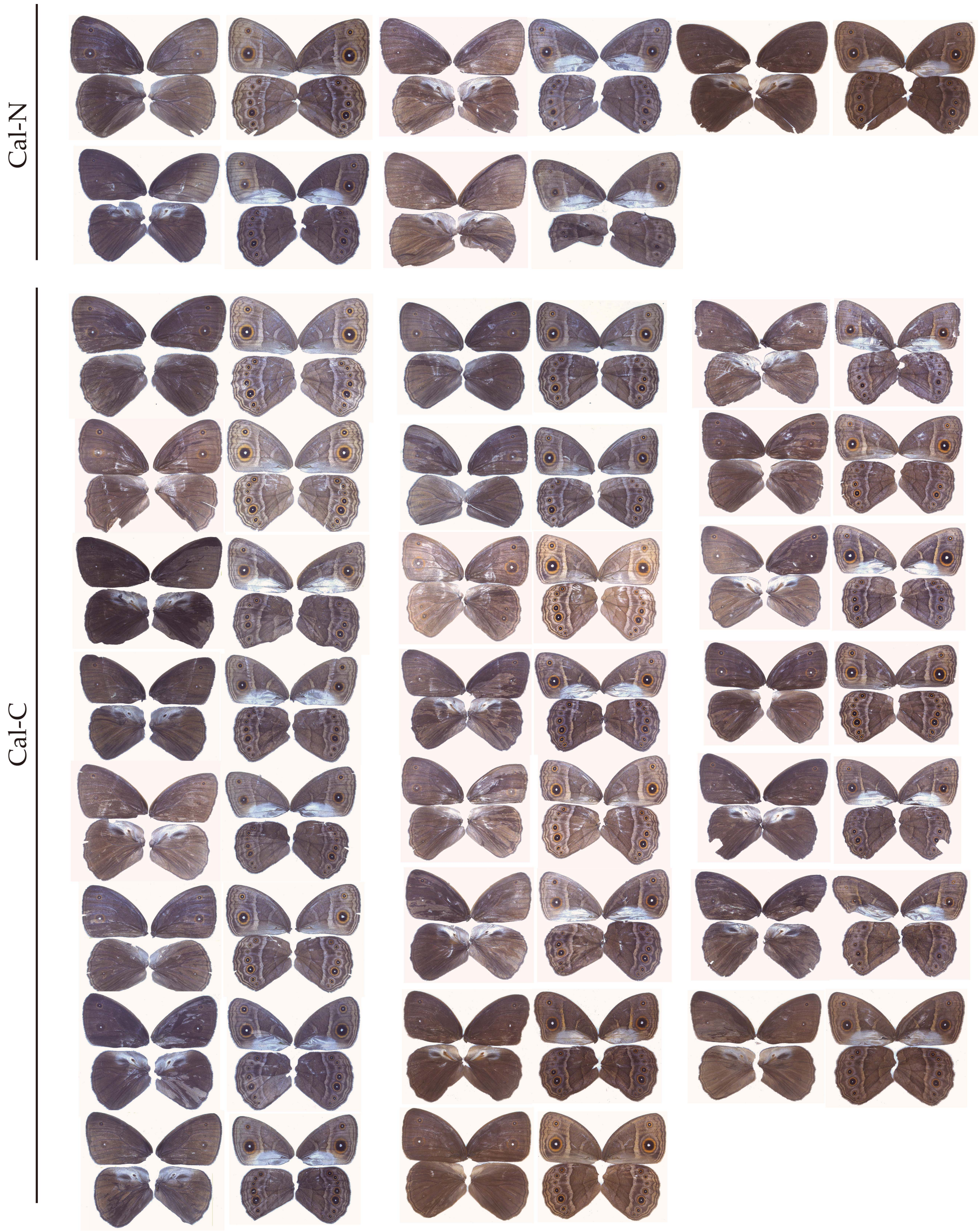


**Figure S2. Calreticulin crispant phenotypes obtained in this study.** Left and right sides of each column show the dorsal and the ventral sides of each individual, respectively.

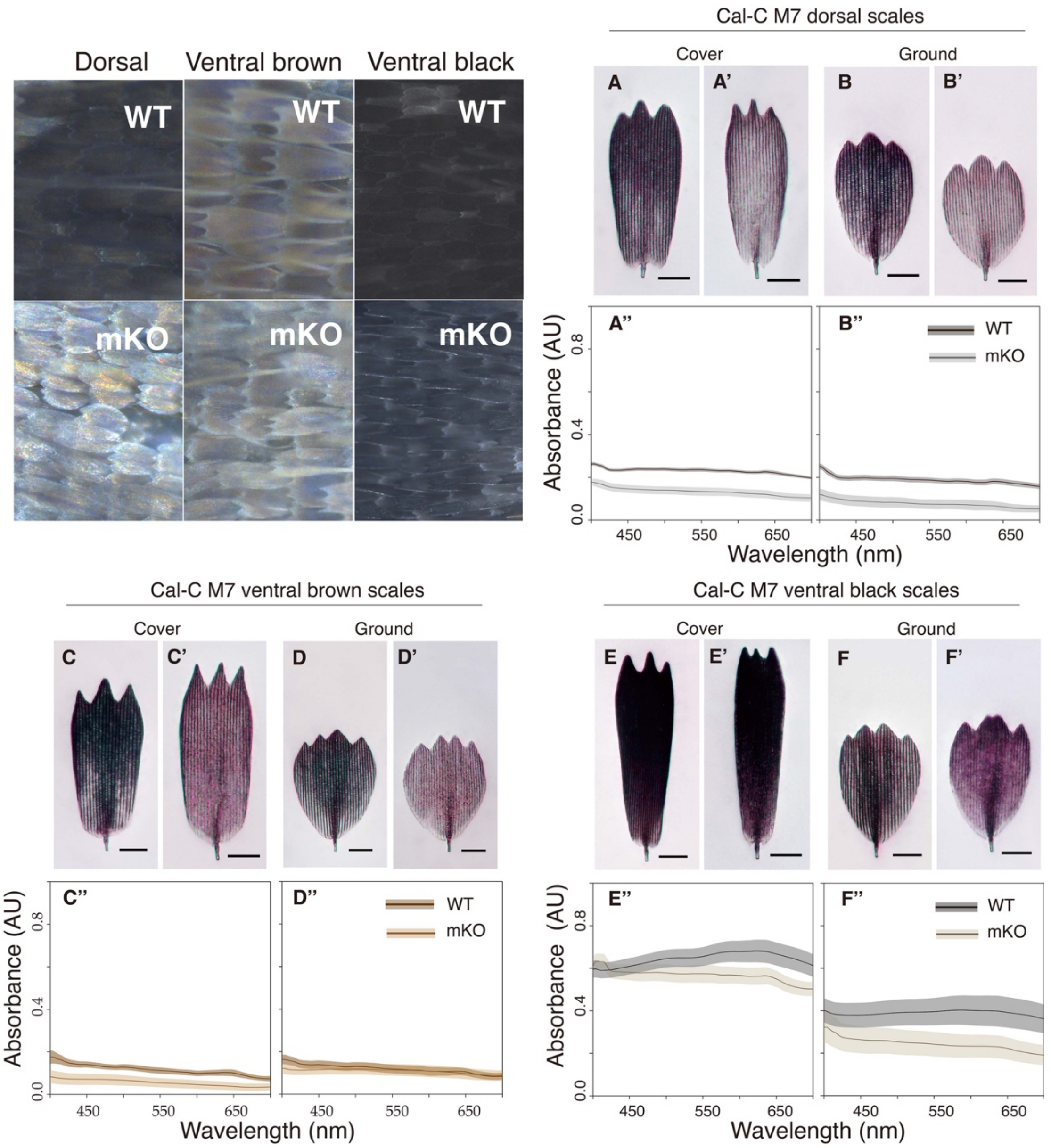


**Figure S3. Disruption of *calreticulin* leads to changes in the color, absorbance spectra, and morphology of brown and black scales.** Optical microscopy images of WT (A–F) and mKO brown and black scales (A'–F') of Cal-C M7. Scale bars: 20 μm. All scales were taken from the same individual. mKO scales show a lighter color, and lower absorbance spectra than WT scales (A''–F''). a.u. = arbitrary units.

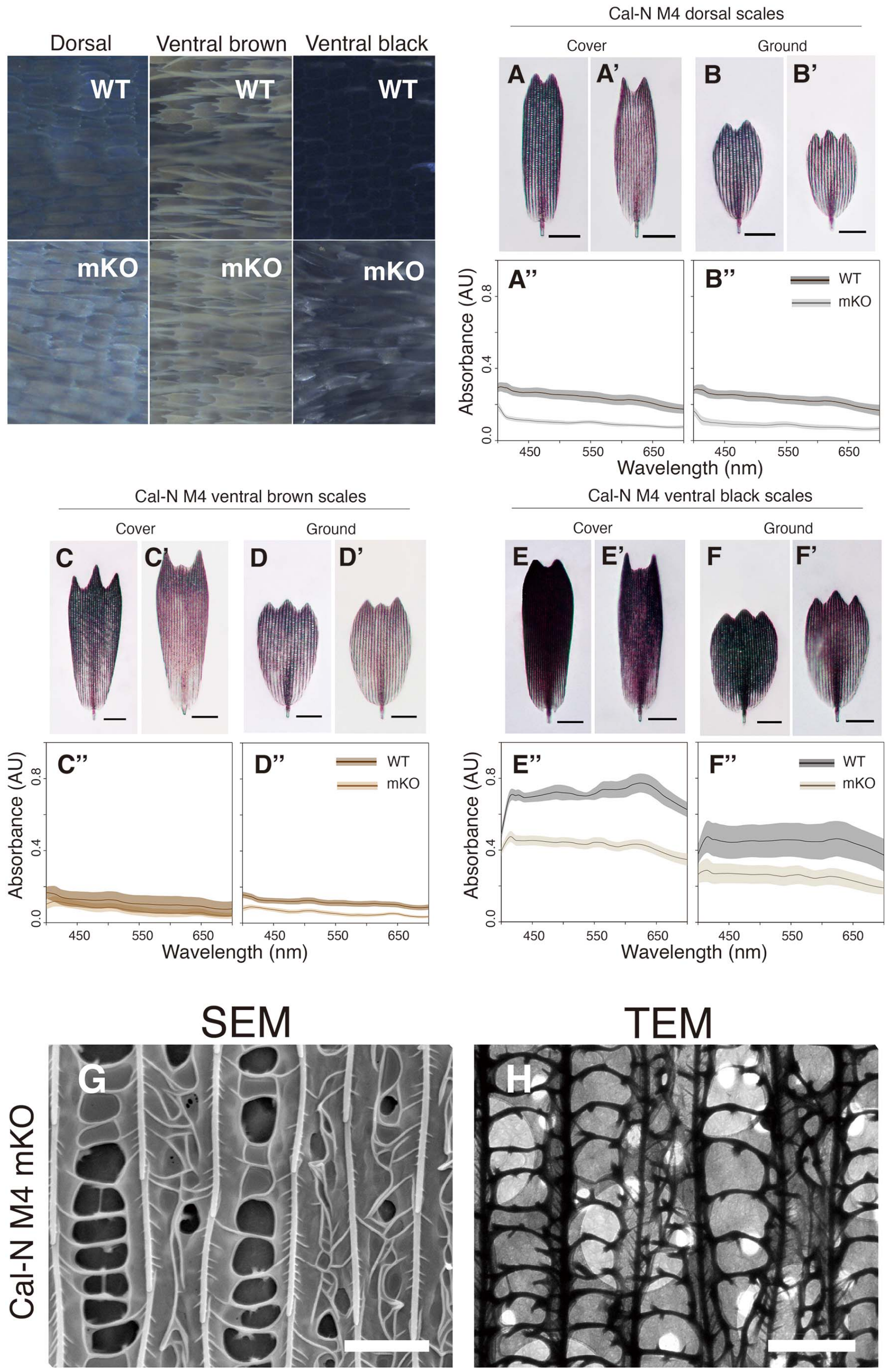


**Figure S4. Cal-N M4 mKO colorful scales show similar absorbance spectra, surface morphology and structure change as Cal-C M7 mKO brown scales.** Optical microscopy images of WT (A–F) and mKO scales (A'–F') of Cal-N M4. Scale bars: 20 μm. All scales are from the same crispant wing. mKO scales show lighter color and blurry cross-ribs. Absorbance spectra of all three-type mKO scales is lower than WT (A''–F''). SEM image (G) and top-down TEM image (H) of the Cal-N M4 mKO dorsal brown scale. Scale bars: 2 μm.

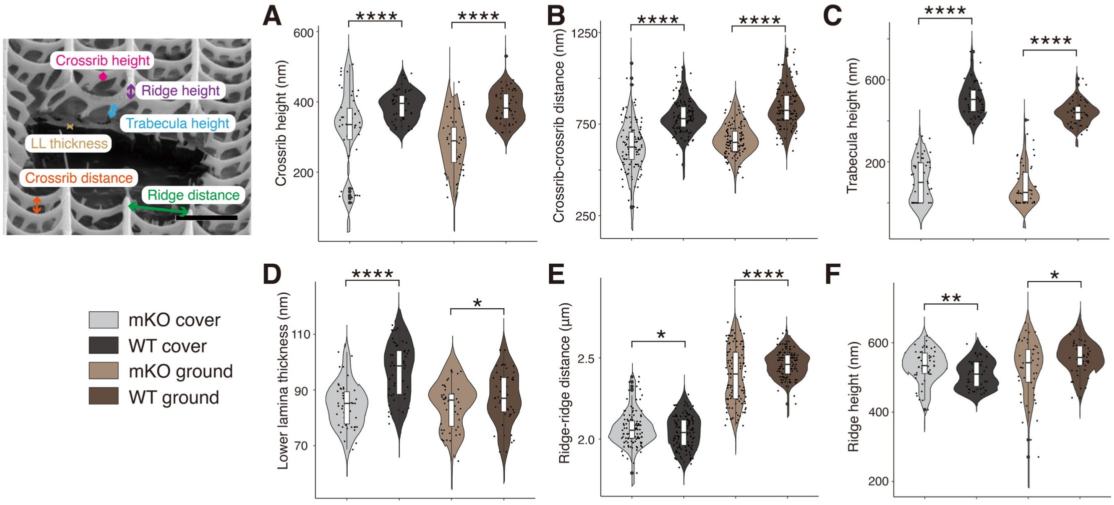


**Figure S5. *calreticulin* knockout leads to shorter trabeculae and shorter cross-ribs that are also more closely spaced, and thinner lower lamina (only in the cover scales).** Violin plots of Cal-C M7 dorsal brown cover and ground scale cross-rib height, n = 5, measurements = 50 (A), adjacent cross-rib distance, n = 5, measurements = 125 (B), trabecula height, n = 5, measurements = 50 (C), lower lamina thickness, n = 5, measurements = 50 (D), ridge-ridge distance, n = 5, measurements = 125 (E) and ridge height, n = 5, measurements = 50 (F). The central line in the violin plot indicates the median of the distribution, while the top and bottom of the box represent the third and first quartiles of the data, respectively. The whiskers show up to 1.5 times the inter-quartile range. *: $p < 0.05$; **: $p < 0.01$; ***: $p < 0.001$; ****: $p < 0.0001$.

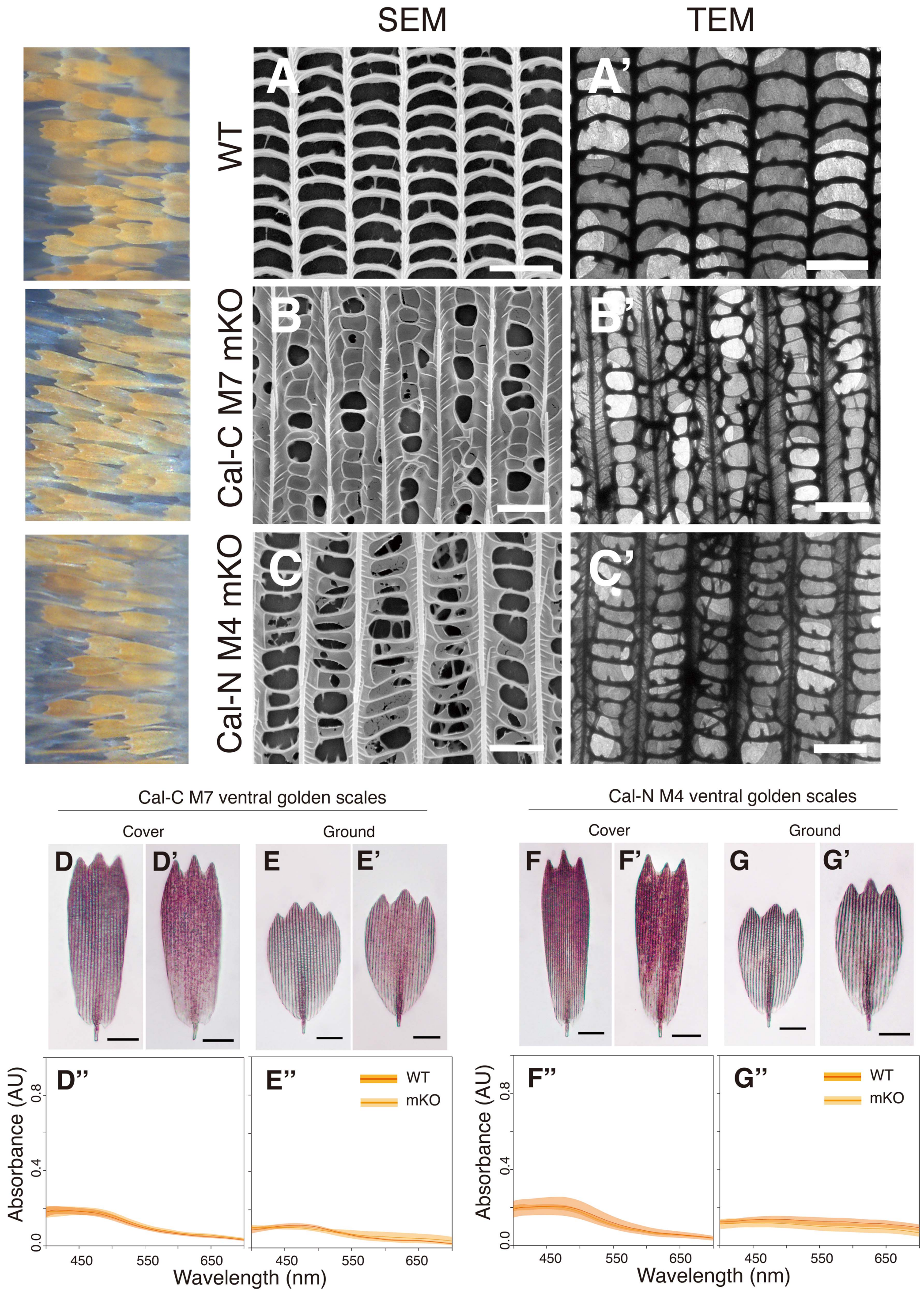


**Figure S6. Calreticulin knockout causes ventral golden scale developmental disruptions, but absorbance spectra were not affected.** (A-C) SEM images of the WT, Cal-C M7, and Cal-N M4 mKO ventral golden scales. (A'-C') Top-down TEM images. Optical microscopy images of WT (D–G), mKO scales (D', E') of Cal-C M7, and mKO scales (F', G') of Cal-N M4. Scales in the same comparison are from the same crispant. The mKO scales from both crispants exhibited the same absorbance spectra as WT (D''–G''). Scale bars: (A-C') 2 µm, (D-G') 20 µm. a.u. = arbitrary units.

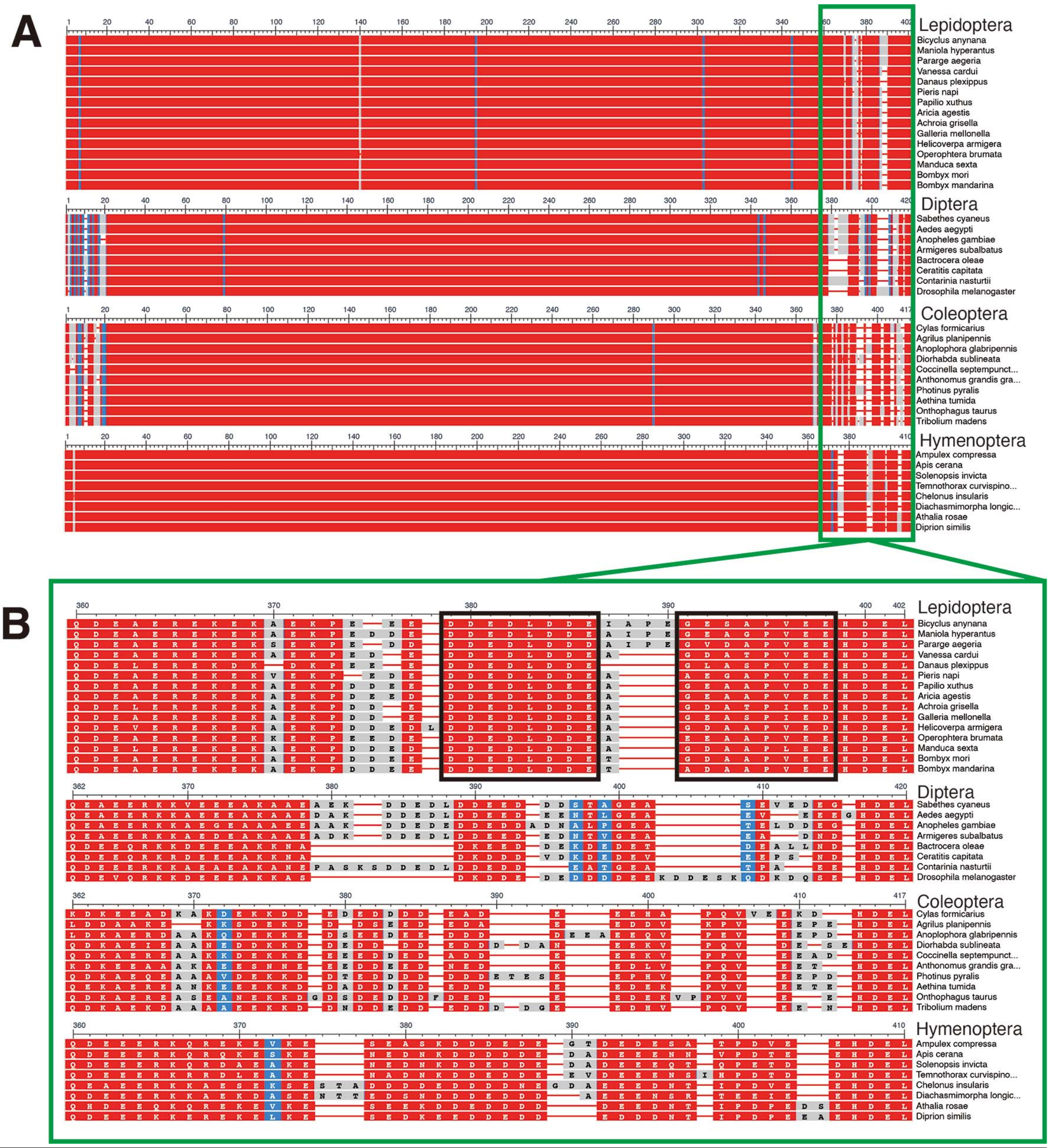


**Figure S7. Two highly conserved motifs are found in the lepidopteran Calreticulin C domain.** Constraint-based Multiple Alignment (COBALT) of representative Calreticulin sequences from four different orders. (A) Whole sequence alignment. (B) The C-terminal ends starting from the conserved Q/L/K site. Black rectangles highlight the two conserved motifs in Lepidoptera. An alignment containing additional species is in Fig. S10 to Fig. S13.

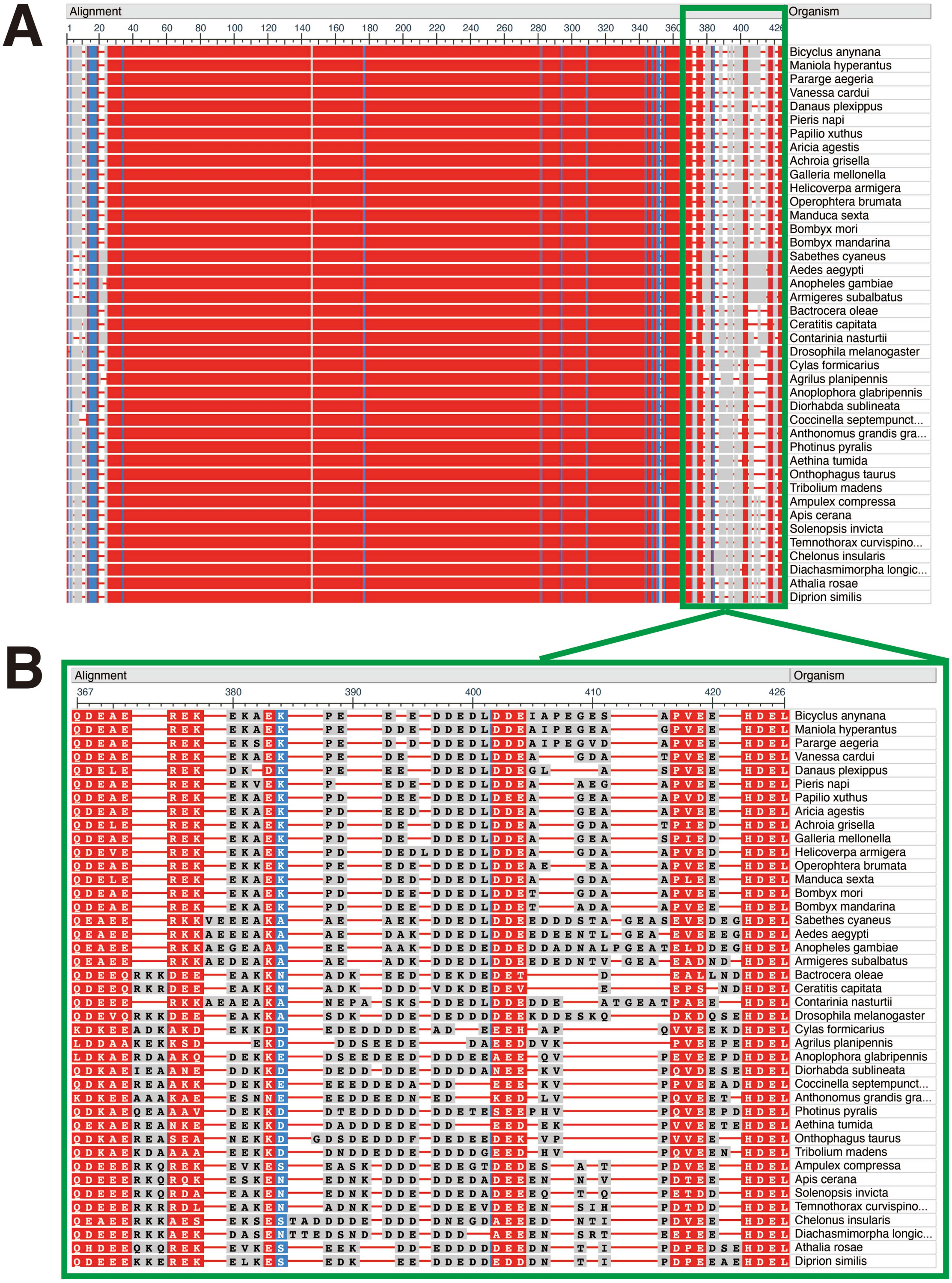


**Figure S8. The conserved C domain motif pattern found in the representative Calreticulin sequences from four different orders is different from lepidopteran.** Constraint-based Multiple Alignment (COBALT) of representative Calreticulin sequences from four different orders in one run. (A) Whole sequence alignment. (B) The C-terminal ends starting from the conserved Q/L/K site. Low sequence similarities indicate this domain varies among insect orders.

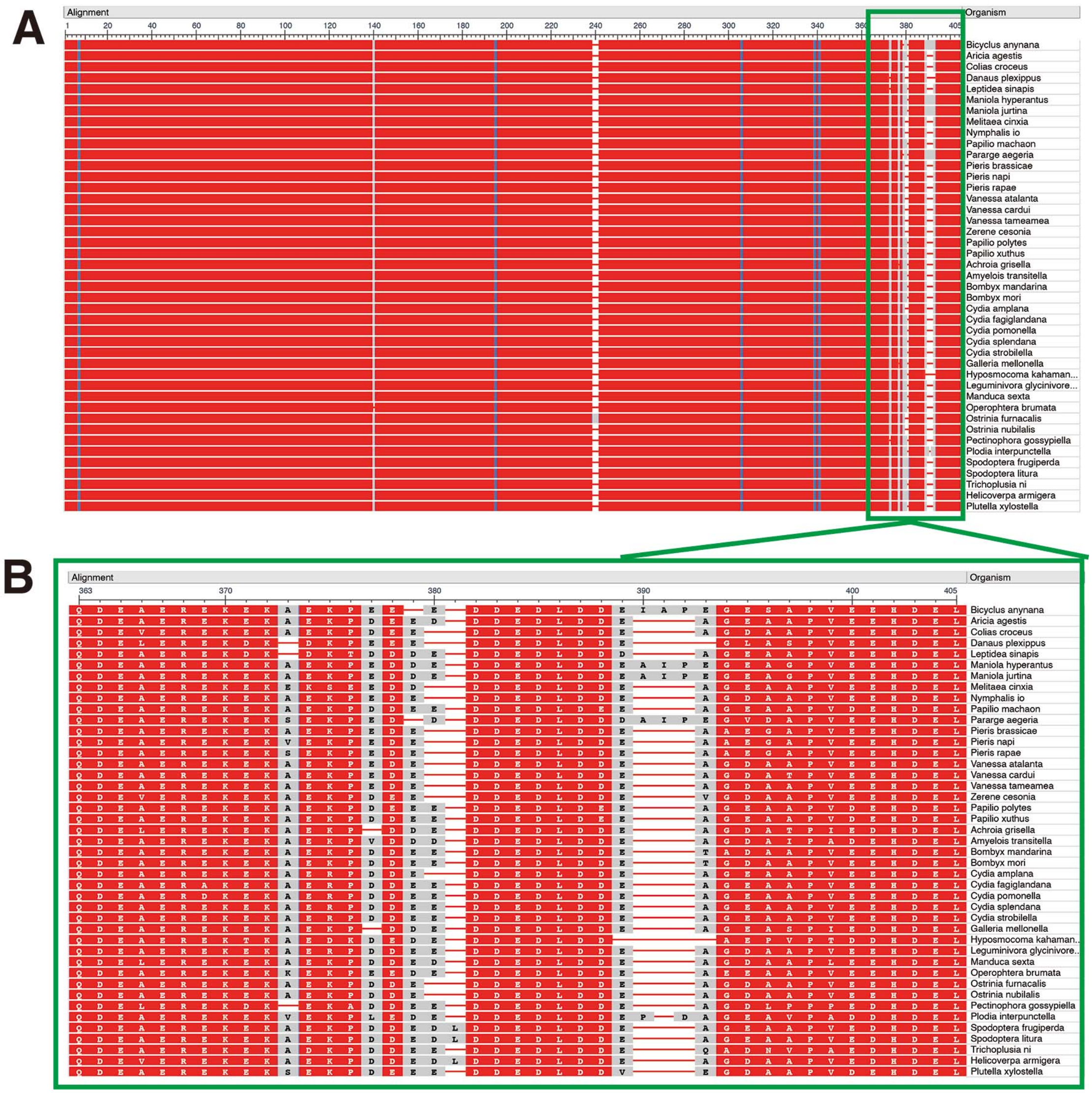


**Figure S9. Constraint-based Multiple Alignment of lepidopteran Calreticulin.** (A) Whole sequence alignment. (B) A zoom-in view of the C-terminal ends starting from the conserved Q/L/K site.

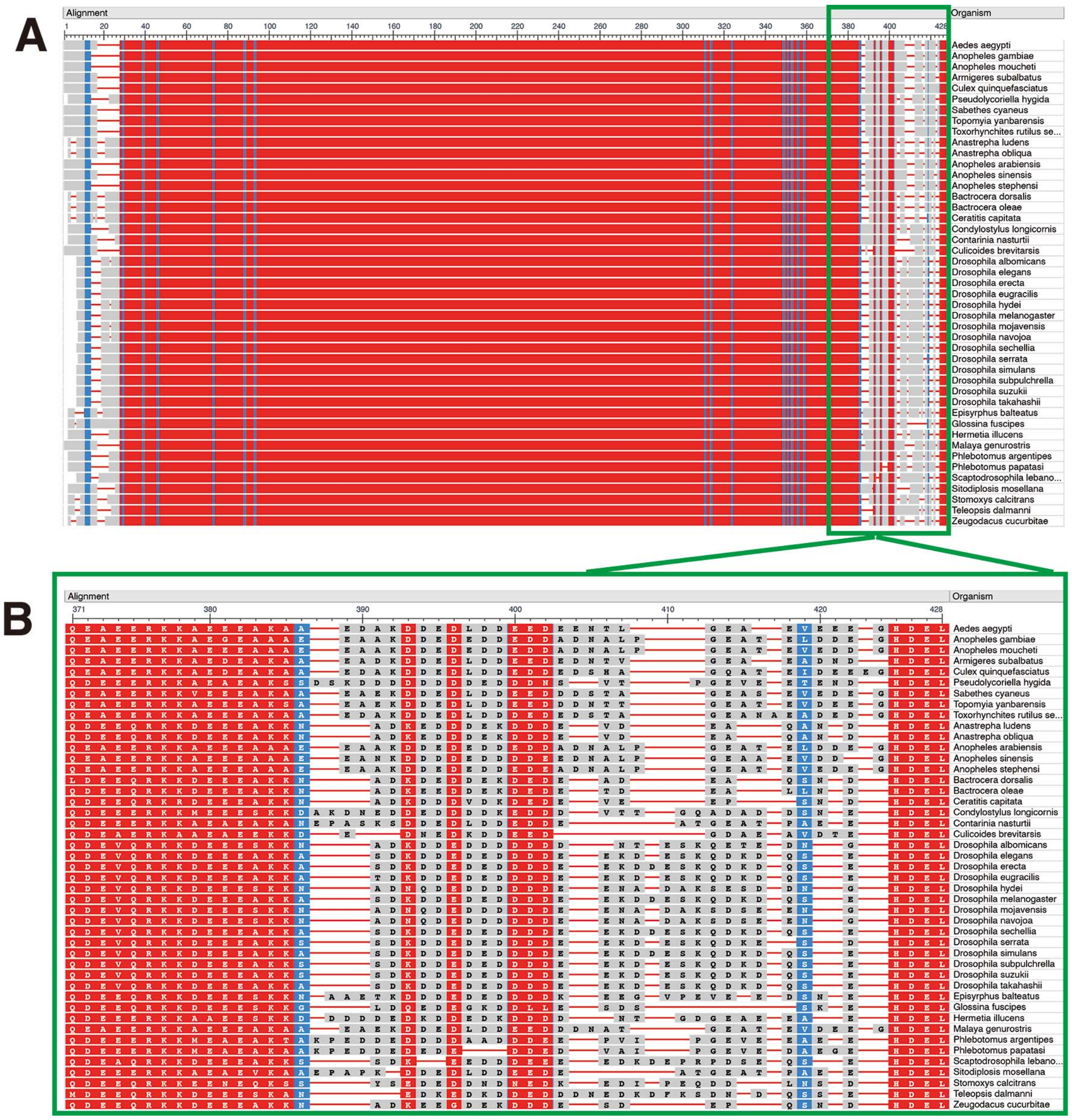


**Figure S10. Constraint-based Multiple Alignment of dipteran Calreticulin.** (A) Whole sequence alignment. (B) A zoom-in view of the C-terminal ends starting from the conserved Q/L/K site.

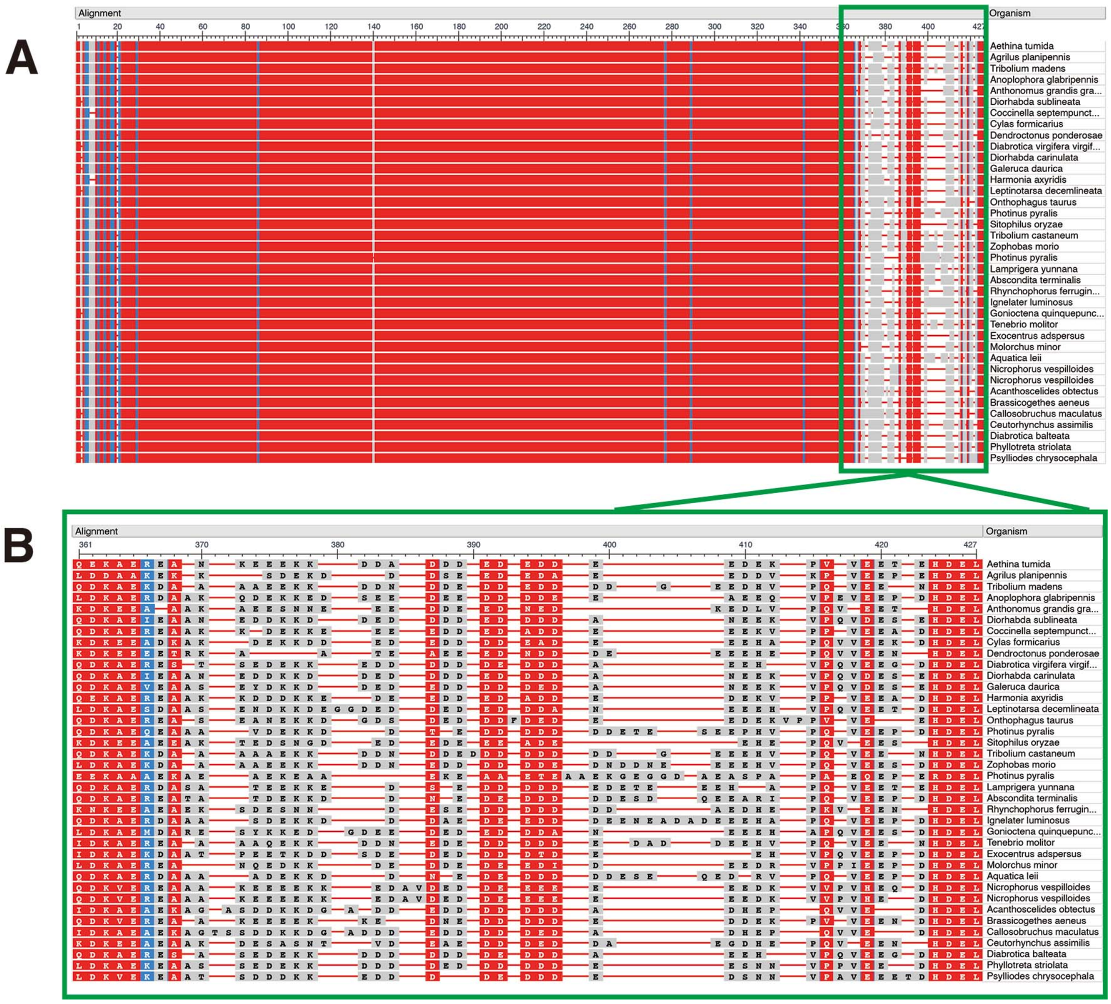

**Figure S11. Constraint-based Multiple Alignment of coleopteran Calreticulin.** (A) Whole sequence alignment. (B) A zoom-in view of the C-terminal ends starting from the conserved Q/L/K site.

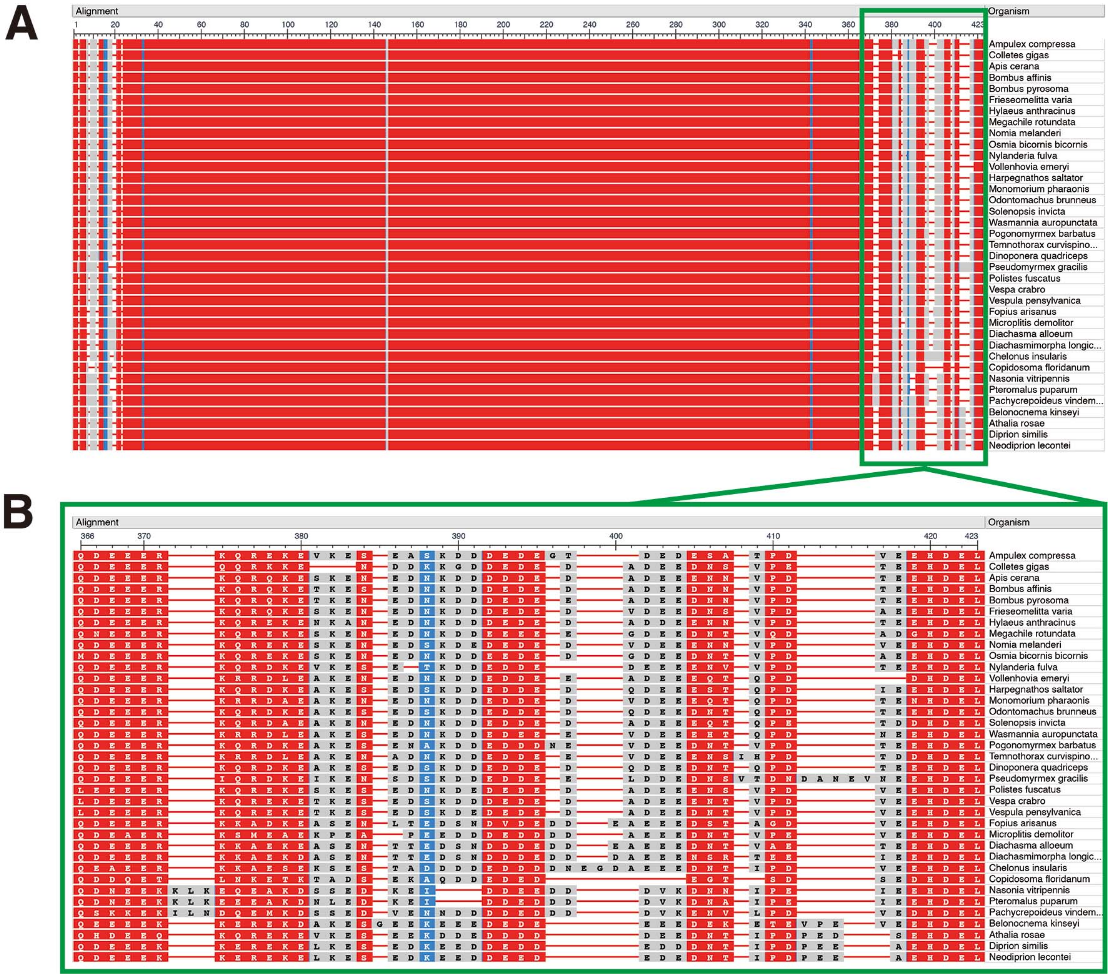


**Figure S12. Constraint-based Multiple Alignment of hymenopteran Calreticulin.** (A) Whole sequence alignment. (B) A zoom-in view of the C-terminal ends starting from the conserved Q/L/K site.
